# Chromosomal Integrons are Genetically and Functionally Isolated Units of Genomes

**DOI:** 10.1101/2023.11.17.567518

**Authors:** Paula Blanco, Filipa Trig da Roza, Laura Toribio-Celestino, Lucía García-Pastor, Niccolò Caselli, Francisco Ojeda, Baptiste Darracq, Ester Vergara, Álvaro San Millán, Ole Skovgaard, Didier Mazel, Céline Loot, José Antonio Escudero

## Abstract

Integrons are genetic elements that increase the evolvability of bacteria by capturing new genes and stockpiling them in arrays. Sedentary chromosomal integrons (SCIs), can be massive and highly stabilized structures encoding hundreds of genes, whose function remains generally unknown. SCIs have co-evolved with the host for aeons and are highly intertwined with their physiology from a mechanistic point of view. But, paradoxically, other aspects, like their variable content and location within the genome, suggest a high genetic and functional independence. In this work, we have explored the connection of SCIs to their host genome using as a model the Superintegron (SI), a 179-cassette long SCI in the genome of *Vibrio cholerae* N16961. We have relocated and deleted the SI using SeqDelTA, a novel method that allows to counteract the strong stabilization conferred by toxin-antitoxin systems within the array. We have characterized in depth the impact in *V. cholerae’s* physiology, measuring fitness, chromosome replication dynamics, persistence, transcriptomics, phenomics and virulence. The deletion of the SI did not produce detectable effects in any condition, proving that -despite millions of years of co-evolution-, SCIs are genetically and functionally isolated units of genomes.

## INTRODUCTION

Integrons are genetic elements that allow bacteria to adapt swiftly to changing environments^1–3^. They recruit new genes encoded in small mobile genetic elements called integron cassettes (IC) (or gene cassettes), stocking them in an array, to form a genetic memory of adaptive functions. According to the needs of their hosts, integrons can also modulate cassette expression by reshuffling their order within the array^4,5^. The integron platform comprises the gene encoding the integrase (*intI*) -the recombinase that governs all reactions-, the integration site where cassettes are incorporated (*attI* site), and two promoters in opposite orientation that drive the expression of the integrase (P_int_) and that of cassettes in the array (P_c_), which are generally promoterless. Integrons are well known for their role in the rise of multidrug resistance and the spread of resistance genes among clinically relevant Gram-negative species^6–8^. Encoded on plasmids, these Mobile Integrons (MIs) act as vehicles for more than 170 resistance genes against the most relevant families of antibiotics^9,10^. Yet, MIs are only a small subset of all integrons that have been mobilized from the chromosomes of environmental bacteria by transposons. The vast majority of integrons in nature are Sedentary Chromosomal Integrons (SCIs), which are found in a variety of phyla and have an ancient origin^11–13^. These SCIs are a highly variable region of the genome, representing a hotspot for genetic diversity^14^. On the antipodes of what is observed for MIs, the functions encoded in sedentary chromosomal integrons are generally not related to antimicrobial resistance and are mostly unknown^15^. The colossal repertoire of cassettes in the environment holds a high potential for biotechnology that is yet to be explored^16^.

The paradigm of SCI is the Superintegron (SI)^17^, a massive structure encoded in the secondary chromosome of *Vibrio cholerae*, the causative agent of cholera disease. In strain N16961, where it was first described, the SI is 126 kb long, embodying 3% of the genome of this bacterium, and containing 179 integron cassettes. Although the model of integron predicts that the SI should be mostly silent, a recent RNAseq analysis shows that many cassettes are expressed at biologically relevant levels (Blanco et al. under review). The functions of cassettes are mostly cryptic, except for the notable exception of 19 cassettes that encode toxin-antitoxin (TA) systems^18,19^ and a chloramphenicol resistance gene^20^. TA cassettes are scattered along the SI and serve to stabilize the array through a post-segregational killing mechanism^21,22^. Despite being the best studied SCI, the knowledge on the Superintegron is limited for two reasons: first, the long cassette array interferes with the genetic tools commonly used in the field to deliver recombination experiments that allow to characterize the system; and second, its integrase cannot be studied in heterologous genetic backgrounds because it seems to require additional host-factors^23–25^.

Sedentary Chromosomal Integrons have co-evolved with the genome of the host for millions of years^26,27^. Indeed, large SCIs like the superintegron are ubiquitous in *Vibrio* species, and the phylogenetic signal of integrases mimics that of the species^1^. This suggests that their acquisition predates the radiation of the genus, an event that occurred more than 300 million years ago^28^. The timescale of this co-evolution process has allowed for the intertwining between host physiology and integron activity at many different levels. A blatant example is that integrase expression is under the control of the host’s SOS response. Yet many other aspects are rather more subtle and represent better the depth of such connections^29^. Examples of this are i) the balanced interplay between the integrase and single strand binding (SSB) proteins of the host on *attC sites*, that regulate *attC* site folding to avoid its structuring unless needed for recombination^30,31^, or ii) the fact that SCIs have a specific orientation related to the origin of replication, to maximize their carrying capacity^22,32^. Yet, a striking observation is that some aspects of SCIs remain extremely plastic, such as their content and their location -despite their size, SCIs can be found in either chromosome depending on the species of *Vibrio*. Hence, it remains unclear whether the intertwining between host and integron is restricted to the functioning of the platform and the mechanistic of recombination (P_int_ and P_c_ promoters and *attC* sites), or if it is extensive to the array of functions encoded in the variable part. In other words, we ignore if cassettes are streamlined to act as plug and play add-ons to the pool of functions encoded in the genome without interacting with it, or whether they can modify or interfere with the host physiology in any other way.

In this work we have addressed the question of whether chromosomal integrons are genetically and functionally isolated units of the genome by relocating and deleting the Superintegron from *V. cholerae*. To overcome the high stabilization of the structure by the TAs, we have designed an approach called SeqDelTA (<u>Seq</u>uential <u>Del</u>etion of <u>T</u>oxin <u>A</u>ntitoxin Systems) that exploits natural competence and homologous recombination to deliver a set of 15 consecutive allelic replacements. At each step, a toxin is inactivated without altering the cognate antitoxin, deleting a piece of the SI. After the last allelic replacement, we produce the scarless deletion of the SI through a counter-selectable suicide vector. The resulting strain has been sequenced, and unintended mutations located elsewhere in the genome have been corrected. Incidentally, this strain represents a milestone in the study of chromosomal integrons, since it provides, for the first time, the possibility of delivering experimental studies using common tools -without interference from the Superintegron. To understand how the deletion of the SI affects *V. cholerae*, we have characterized in depth this strain from a variety of perspectives, including chromosome replication dynamics, fitness, persistence, transcriptomics, phenomics and virulence assays. We observe no significant variations in the physiology of the ΔSI strain compared to the wild type. We conclude that sedentary chromosomal integrons are functionally isolated units of the genome, despite eons of co-evolution with their hosts. This has strong implications in the type of functions one can expect to find in integron cassettes.

## RESULTS

### Genomic position of the SI does not alter *V. cholerae* growth

The genomic location of SCIs in Vibrio species is remarkably plastic given their size. We ignore whether SCI relocations in different species have been innocuous for the cell or have entailed complex changes that required subsequent adaptation. To test this, here we used the Φ HK recombination machinery to relocate the SI. We moved it from the right arm of chromosome 2 -close to the replication terminus-, to chromosome 1, near the origin of replication, where we placed it in both orientations. This relocation could have a potential impact in cell growth because of the higher gene dosage effect near the Ori of chromosome 1. Additionally, it could also generate conflicts between transcription and translation machineries depending on the orientation of the array. We measured growth of both mutants and could not observe differences with the WT, suggesting that SCIs are genetically independent units of the genome.

**Figure 1.**
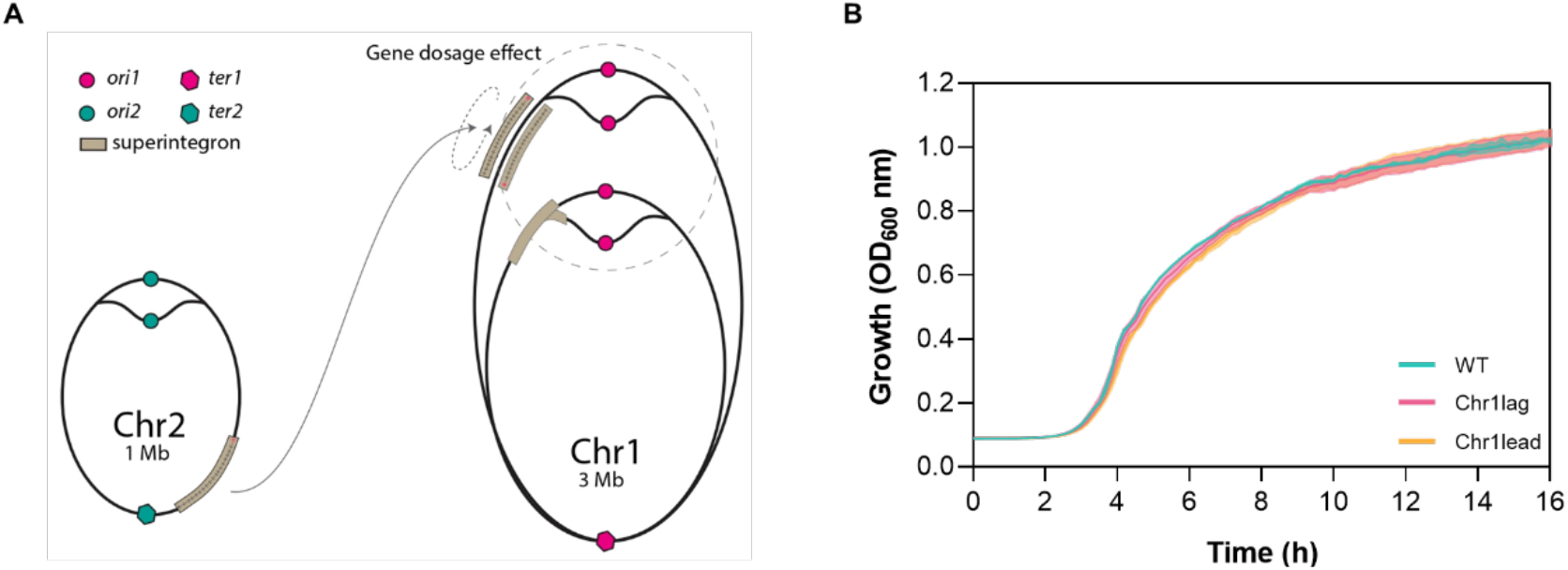
(**A**) Scheme of the relocation of the SI from Chr2 to Chr1, where it is inserted in both orientations (modified from^33^). (**B**) Growth curves of WT and relocated-SI strains (Chr1lag and Chr1lead). No significant growth differences are observed.

### SeqDelTA, a tool to erase chromosomal integrons

To further assess the level of genetic and functional independence, we sought to delete the superintegron. The deletion of the SI in *V. cholerae* is a longstanding milestone in the field, that should allow to deliver the kind of experiments that have advanced the field in the last years using the Class 1 integron as a model. Previous efforts to delete the SI were hampered by unknown TA systems and by a one-step deletion approach. The recent full characterization of TA systems in the SI^18,19^ has enabled the development of SeqDelTA, a multi-step approach that allows to sequentially inactivate TA systems while deleting, at each step, the cassette cargo between two TAs (Figure 2).

**Figure 2.**
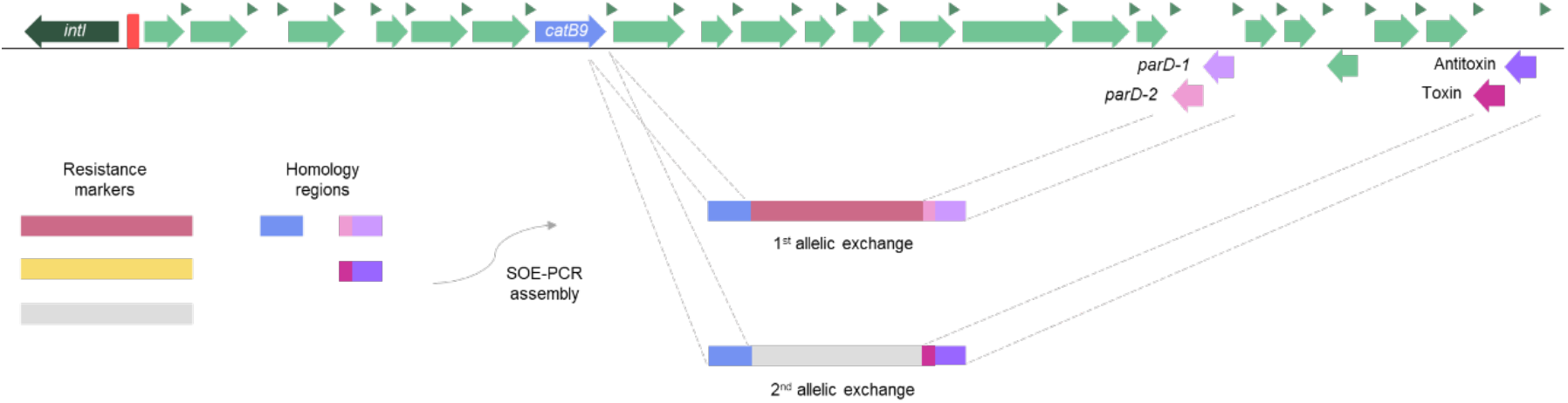
Schematic representation of two successive allelic replacements using SeqDelTA. The left homology region is conserved during the deletion process, allowing to recycle the resistance markers at each step. The right homology region is redesigned at each step to sequentially inactivate the TA modules. This allows to inactivate a TA system by knocking the toxin but leaving the antitoxin intact. A total of 18 sequential replacements were performed. The last deletion step was carried out using a suicide plasmid.

Briefly, we take advantage of *V. cholerae*’s natural competence to provide linear DNA fragments in which a resistance marker is flanked by left and right homology regions (LHR and RHR). These HRs contain the sequence of specific parts of the superintegron to produce allelic replacements. The superintegron spans from gene VCA0291 -the integrase-to VCA0506 (nucleotides 309,750 to 435,034). In the design of the allelic exchanges, we used a conserved LHR targeting the *catB9* chloramphenicol resistance cassette in 9^th^ position of the array (VCA0300). The rationale for this choice is that, since the strain is susceptible to chloramphenicol, this region of the SI is expected to be silent, being therefore a good starting point in the 5’ end of the array, free of interference with the P_c_ promoter. In contrast, the RHR changes at each step of the deletion, targeting the next TA downstream the array. RHRs were designed to produce crossovers that maintained the antitoxin of each system intact while deleting the toxin gene. Keeping a common LHR while advancing the deletion through changes in RHRs allowed the removal of the resistance marker introduced in the previous step, easily alternating resistance markers. This scheme for SeqDelTA was generally kept constant. In the cases in which the orientation and/or the order of the genes in the TA system where not favorable, we used alternative LHRs and maintained two resistance markers in consecutive allelic replacements. Then, a third replacement allowed to remove both markers. After each allelic exchange, several colonies growing on the appropriate antibiotic were verified through PCR and checked phenotypically for the loss of the previous marker where applicable. The growth curve of PCR-verified clones was then analyzed to avoid the hitchhiking of mutations with important deleterious effects. A clone without growth defects was chosen to continue the deletion process. After the last allelic replacement, we had erased all cassettes from VCA0300 (*catB9*) to VCA0503 (the last toxin). We then used a counter-selectable integrative vector (pMP7^34^) to deliver the scarless deletion of the remainder of the SI -including the integrase, the first 8 and the last 2 cassettes from the array, as well as a transposase inserted immediately downstream the last cassette (VCA0291 to VCA0508). A PCR from the borders of the integron confirmed the deletion of the SI.

### Genome Sequencing and Marker Frequency Analysis

We performed genome sequencing on the obtained *V. cholerae* ΔSI using Illumina short reads and confirmed the scarless deletion of the SI. We also used this approach to perform Marker Frequency Analysis (MFA), a technique that allows to assess the dynamics of chromosome replication. MFA compares the abundance of reads across the genome between exponential and stationary growth conditions. In fast growing bacteria as *V. cholerae*, this comparison reveals the gene dosage gradient from the origin to the terminus of replication as an inverted “V”. MFA allows to observe the relative timing of replication of both chromosomes, and the speed of replication across the genome^35^. As shown in Figure 3, no major alterations in chromosome replication dynamics could be observed, compared to the WT strain.

**Figure 3.**
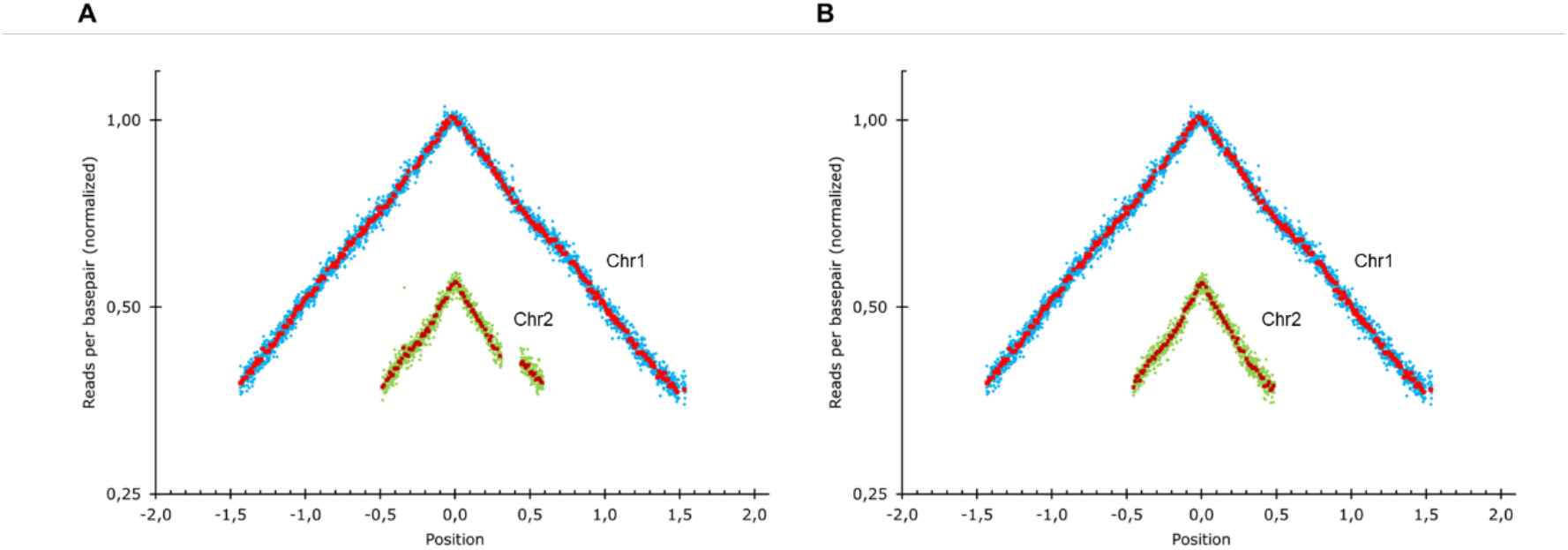
(**A**) Marker frequency analysis (MFA) of the two chromosomes of *V. cholerae* ΔSI mapped against *V. cholerae* WT genome and (**B**) against *V. cholerae* ΔSI genome. The gap on chromosome 2 in panel A reflects the superintegrons deletion. The genome position is represented relative to the *oriC* (set to 0). Blue and green rhombus represent the average of 1 kbp windows, and red and garnet squares the average of 10 kbp windows in Chr1 and Chr 2, respectively.

MFA can reveal the presence of large chromosomal rearrangements (such as inversions), through inconsistencies in the slope of the inverted V shape, but lacks resolution for smaller rearrangements like IS movements. We therefore sequenced *V. cholerae* ΔSI using MinION long reads to detect the presence of any rearrangement that could go undetected by the assembly of short reads. Combination of small and long reads allowed to obtain a high-quality genome sequence in which we could rule out genomic rearrangements while detecting the presence of unintended single nucleotide polymorphisms (SNPs). The *V. cholerae* ΔSI strain bore three unintended mutations in *rocS* (VC0653), *cry2* (VC01392) and *rpoS* (VC0534) genes. RocS is a diguanylate cyclase involved in the smooth-to-rugose switch in our strain. A101 had a single base deletion that altered the reading frame of *rocS*. We could observe the loss of motility and increase in biofilm formation described in *rocS^-^* mutants^36,37^. We were able to map the emergence of this mutation to the third deletion step of SeqDelTA. Interestingly, 12 clones from two independent experiments had been kept from this deletion step, of which 11 had mutations in *rocS*. We found 8 different types of mutations among them, pointing to an unusually high mutation rate. Indeed, the sequence of *rocS* is rich in homopolymeric tracts, and is likely to act as a contingency locus, allowing for a frequent smooth-to-rugose switch^38^. *cry2* presented a non-synonymous (Pro129Lys) mutation. This gene encodes a protein annotated as a photolyase, for which no experimental evidence of its function could be found. RpoS is an alternative sigma factor related to stress response and entry in stationary phase. It is a major regulator of cell physiology and influences many relevant phenotypes, such as oxidative stress, natural competence, motility, or colonization^39–41^. *V. cholerae* ΔSI harbored a single-base insertion in *rpoS* giving rise to a frameshift.

To understand the impact of the deletion of the SI in the bacterium’s physiology we need to avoid the interference of the mutations in these three genes. To do so, we restored the WT sequence of the three alleles using the counterselectable suicide vector pMP7. We then verified the restoration of the three genes, as well as the absence of other mutations arising during this process again through whole genome sequencing using short and long reads. As a result, we could confirm that the ΔSI strain is genetically identical to the WT except for the absence of the SI and the loss of one of the two copies of satellite phage RS1.

### Transcriptomics

Regulatory networks are generally finely tuned and often affected by minor genetic changes. To address the level of functional isolation of the SI, we sought to assess the changes in expression patterns of the genome in the absence of the SI. Indeed, the deletion of the SI could have pleiotropic effects through direct and indirect links to regulatory networks within the cell. To unveil such effects, we have performed transcriptomic analysis of the ΔSI and the parental strain in exponential and stationary phase. As expected, all genes contained within the SI gave a clear signal of decreased expression. Unexpectedly, the transcriptome of the rest of the genome remained fundamentally unaltered, with few exceptions that were confirmed through RT-qPCR using independent RNA extractions (Figure 4A and 4B).

**Figure 4.**
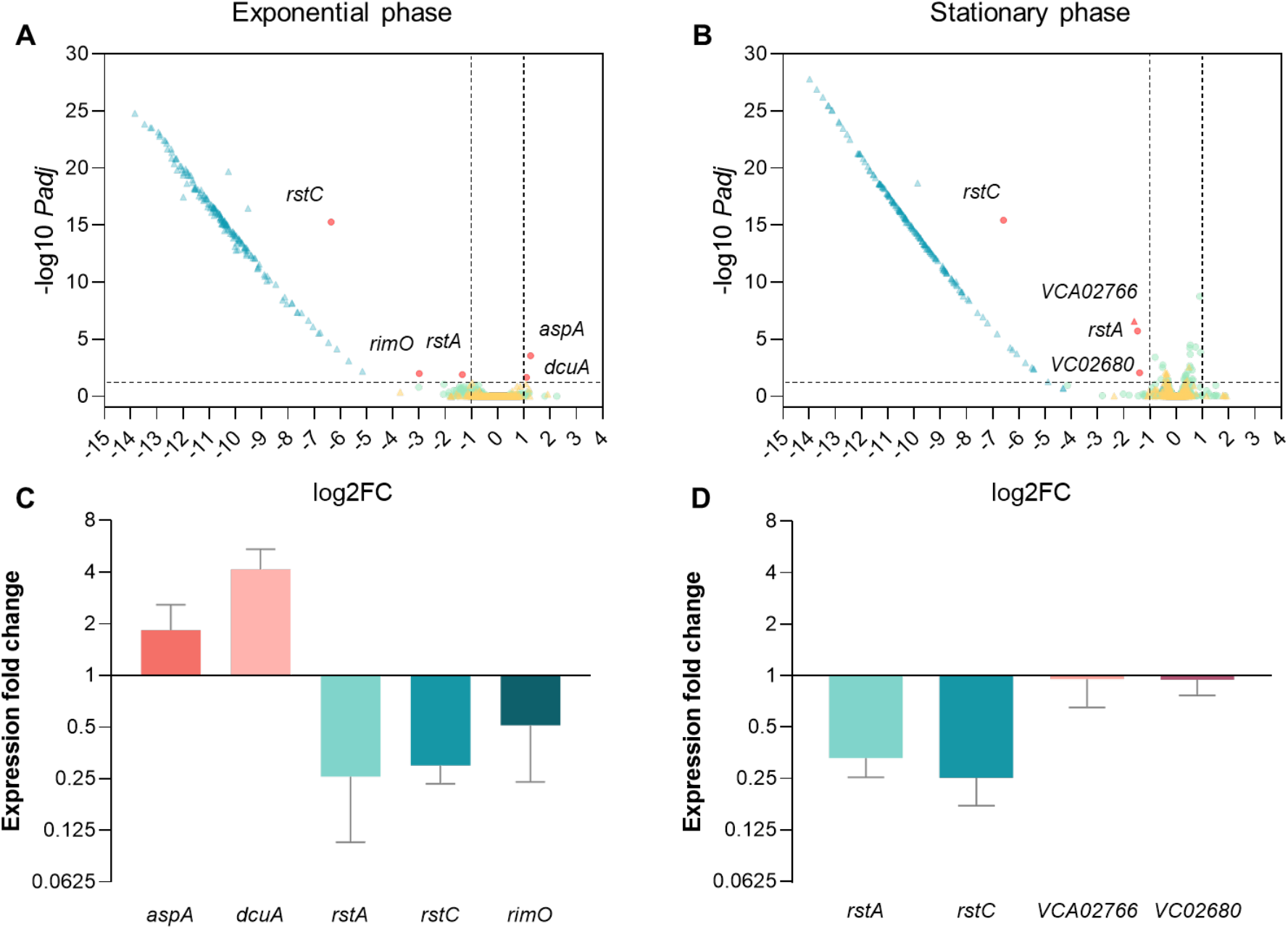
(**A**) Volcano plots showing gene expression measured by RNA-seq in *V. cholerae* ΔSI in comparison to the WT strain in exponential and (**B**) stationary growth phase. Green dots represent chromosome 1 genes; yellow triangles represent chromosome 2 genes, while blue triangles represent those genes from the superintegron; red dots or triangles represent downregulated or upregulated genes from chromosome 1 or 2, respectively. The vertical dashed lines indicate the log_2_ fold change (FC) cutoffs, and the horizontal dashed line indicates the threshold of the *Padj* value (< 0,05). (**C**) Validation of the differentially expressed genes by quantitative PCR (qPCR) during exponential and (**D**) stationary growth phase. Differential expression values represent the fold change in gene expression compared to the WT strain. Error bars indicate standard deviation of two biological replicates with three technical replicates each.

The loss of one of the tandem copies of RS1 in the ΔSI strain led to the downregulation in both exponential growth and stationary phase of *rstA* (VC01411) and *rstC* (VC01409), that was confirmed through RT-qPCR on independent samples. In the WT strain, *rstA* is found three times as part of the Φ CTX (VC01397) and two tandem copies of the satellite phage Φ RS1 (VC01407, VC01411), while *rstC* is exclusively found in the latter in two copies (VC01409, VC01413). ^394041^ This downregulation trough the loss of a Φ RS1 copy serves as an additional validation of transcriptomic results.

Besides prophage-related genes, in exponential growth phase we also found a two-fold upregulation of *aspA* (VC00205; aspartate ammonia-lyase) and *dcuA* (VC00204; C4-dicarboxilase transporter) and a 6-fold downregulation of *rimO* (VC00523; ribosomal protein S12 methylthiotransferase). *aspA* and *dcuA* colocalize in *V. cholerae* chromosome 1. While *aspA* catalyzes the reversible conversion of L-aspartate to fumarate^42^, *dcuA* is involved in the L-aspartate/fumarate antiport^43^. Aditionally, RNAseq data showed genes VC02680 and VCA02766 to be down-regulated 2-fold approximately in stationary phase, but RT-qPCR failed to confirm such downregulation (Figure 4C and 4D).

A gene set enrichment analysis (GSEA) can reveal the collective up- or downregulation of related genes, even if individually they show no significant changes in expression patterns. GSEA was performed here on the group of differentially expressed genes, revealing that the ΔSI strain showed a mild enrichment of the iron ion transmembrane transport process (GO:0034755; NES = 2,126432; *Padj* = 0,00714409) and the arginine biosynthetic process (GO:0006526; NES = 1,959273; *Padj* = 0,02539212) in exponential phase and of isoleucine biosynthetic process in stationary phase (GO: 0009097; NES = 2,020381; *Padj* = 0,0129091). Collectively, these data indicate that deleting the superintegron does not have important consequences at the transcriptome level in *V. cholerae*.

### Phenotype search

The working model of integrons suggested that most of the array of the superintegron is silent, and only the cassettes closest to the P_c_ are being expressed. Hence, it would be unlikely to find many phenotypic changes after the deletion of the SI. Yet this seems not to be true, with independent reports showing both the presence of transcription start sites scattered along the superintegron, and a large proportion of cassettes expressed at biologically relevant levels (Blanco *et al.* under review.), in accordance with our RNAseq results^44^. To address the potential emergence of phenotypic changes, and therefore the discovery of cassette functions, we implemented a broad approach to characterize a maximum of phenotypes.

### Growth and Fitness

Deletion of *V. cholerae* superintegron implies the loss of the 3% of the bacterial genome. Thus, it is expected that elimination of this genomic load would result in some growth and/or fitness consequences. To explore this possibility, we performed growth curves of 24 independent *V. cholerae* ΔSI colonies and compared with the WT in rich LB medium. As shown in Figure 5A, growth curves of both strains present similar shapes after 16 h of growth. We did not observe significant growth differences between strains measured as Vmax (Figure 5B). We then measured the area under de curve (AUC), which combines more information about the growth curve, such as lag phase, Vmax and the carrying capacity. The results showed similar AUC values for both strains (Figure 5C). To confirm that the deletion of the superintegron does not influence bacterial fitness, we increased the sensitivity by performing competition assays using flow cytometry. Using a fluorescent *E. coli* strain as competitor, we found no significant differences between the fitness values given by *V. cholerae* WT and ΔSI. Altogether, these results indicate that deletion of the *V. cholerae* superintegron does not affect significantly bacterial growth.

**Figure 5.**
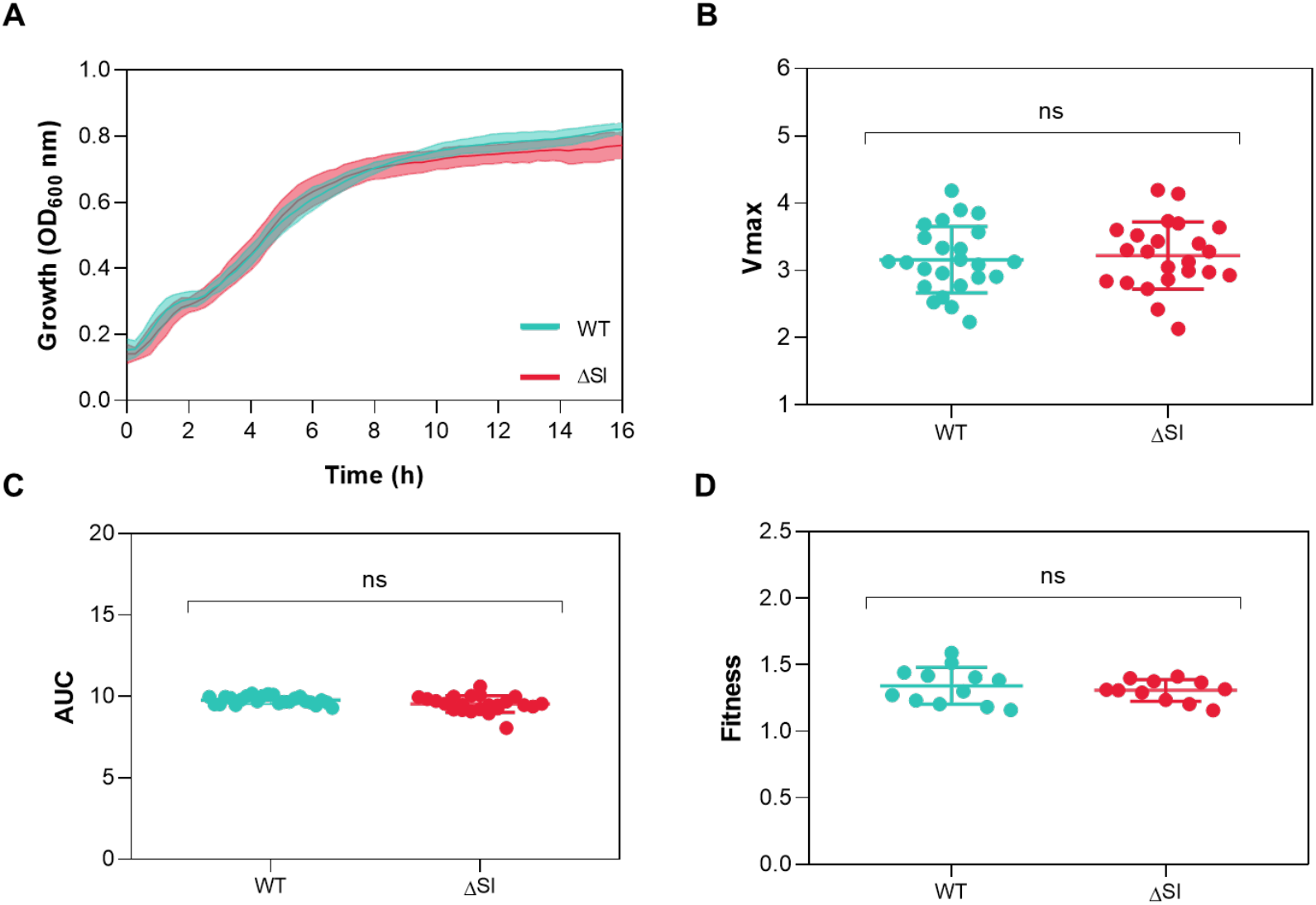
(**A**) Growth curves of *V. cholerae* WT (blue) and ΔSI (red) strains in LB. Growth parameters including (**B**) Vmax and (**C**) the area under the curve (AUC) were extracted from growth curves in LB by measuring the optical density at 600 nm (OD_600_). Values correspond to the measurement of 24 independent colonies. (**D**) Relative fitness of *V. cholerae* WT and ΔSI strains compared with *E. coli* DH5α PcS::*gfp*. Competition assays were performed in LB by inoculating a cells ratio of 1:1. Fitness values were determined from 12 independent experiments by flow cytometry. The p-values were calculated by comparing each measure with that of the WT strain using unpaired t-test. Ns: not significant.

### Biolog Phenotype Microarrays

To address if we can observe phenotypic differences in the absence of the SI, we have used Biolog Phenotype Microarrays® (PMs), a phenomics technology that allows the testing of bacterial respiration and growth in nearly 2.000 conditions^45^. PMs test carbon utilization, nitrogen sources, phosphorus and sulfur compounds, biosynthetic pathways, osmotic, ionic and pH effects, as well as a broad set of chemicals and antibiotic compounds at four different concentrations. By using tetrazolium violet as a redox dye in the mixture, we monitor bacterial respiration for 24 hours and compare the metabolism of two independent replicates of the WT and ΔSI strain in all the available conditions (Supplementary Figure 1).

To search for differences between the two strains, the mean AUCs given by both replicates of WT and ΔSI in a specific compound is graphically compared using Matlab software (R2022a). In Figure 6, we show the correlation of AUC values for WT and ΔSI strains in each condition. Distance to the origin of the axis represents metabolic activity, and deviations from the diagonal represents differences between strains. Generally, dots fall within the diagonal, showing that both strains behave similarly in the presence of the different compounds, which also supports the similarity in growth rates across a broad variety of conditions. It is of note, that the ΔSI shows a potential decrease in respiration when using dipeptides and tripeptides as nitrogen sources (Plates 6 and 8) (see discussion). Within antimicrobial-containing plates (PM11 to 20), we generally did not found differences between the WT and ΔSI. However, we have highlighted some isolated cases in Figure 6 where the AUC of one of the strains is at least the double of the other and the standard deviation does not cross the bisector. These compounds are: ceftriaxone (G4, PM11), cadmium chloride (D4, PM14), methyl viologen (E12, PM15), norfloxacin (B3, PM16), tannic acid (F10, PM17), sodium metasilicate (E4, PM18), iodonitro tetrazolium violet (D8, PM19), thioridazine (C1, PM20), and benserazide (A10, PM20). Despite the lack of consistent results with other molecules of the same family, we believed these results deserved further verification. We selected three compounds (ceftriaxone, norfloxacin and tannic acid) and measured the MIC, but found identical values for both strains: norfloxacin 50 ng/ml; ceftriaxone 25 ng/ml; tannic acid 250 ng/ml. We also performed growth curves using subinhibitory concentrations of the three compounds in MH, but could not reproduce the results obtained using phenotype microarrays (Figure 7), suggesting that phenotypes.

**Figure 6.**
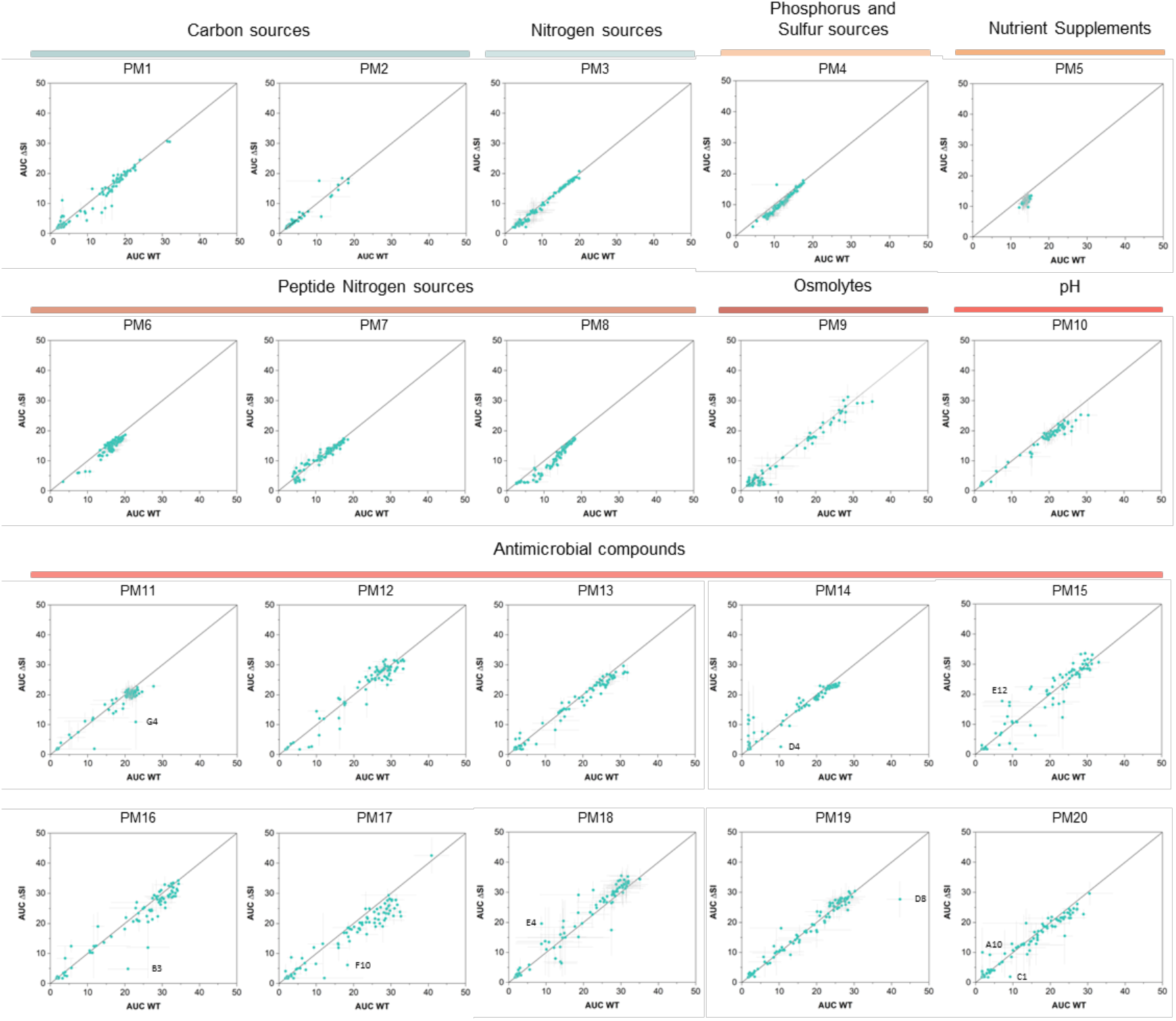
Comparison of the area under the curves (AUC) values of *V. cholerae* WT and ΔSI in different substrates given by Biolog Phenotype Microarrays (PM1-PM20). The AUC values are the mean of two biological replicates for each strain. Error bars indicate the standard error. The points that are above the bisector indicate those conditions where the absence of the superintegron is beneficial for the bacteria, while those points that lay under the bisector indicate the conditions where the presence of the superintegron is beneficial for the bacteria.

**Figure 7.**
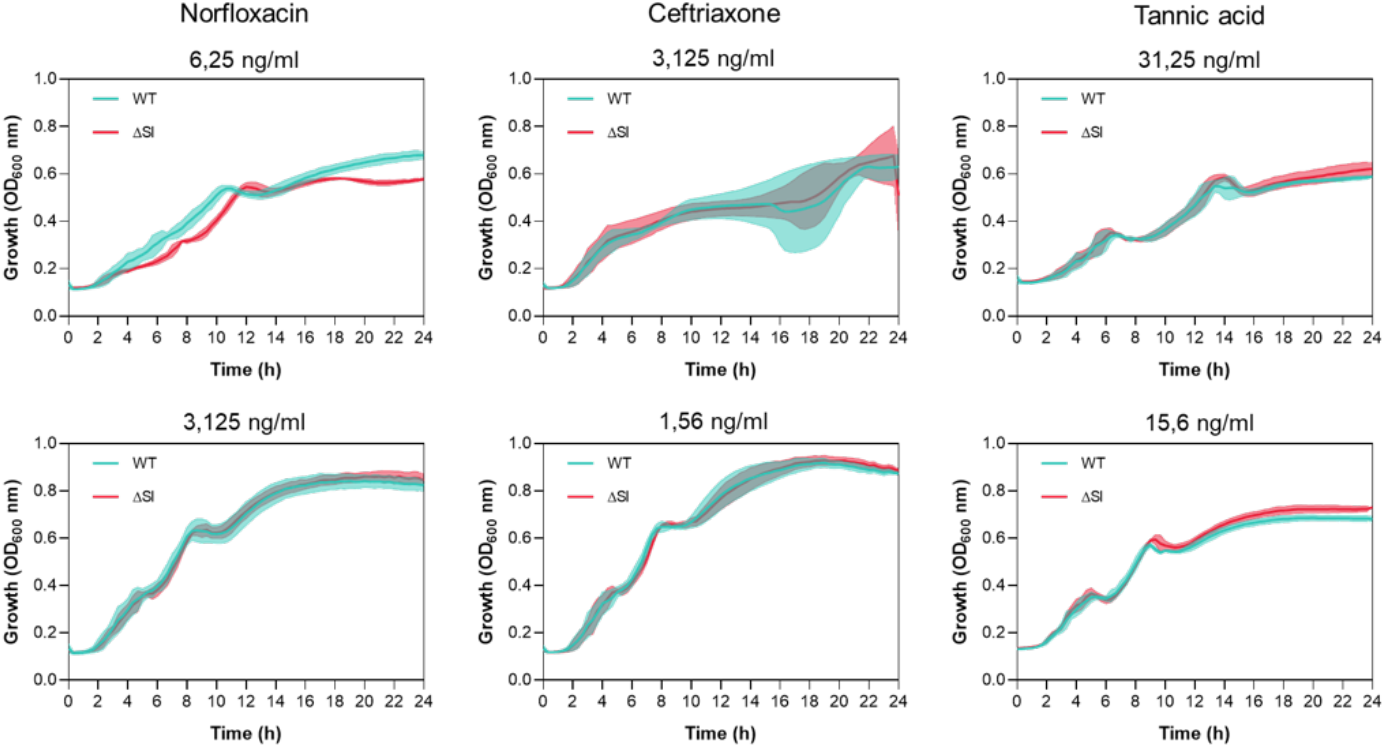
Growth curves of *V. cholerae* WT (blue) and ΔSI (red) strains in MH medium containing subinhibitory concentrations of norfloxacin, ceftriaxone or tannic acid. Colored shading of the curves represents the standard deviation of at least three biological replicates.

### Persistence, virulence, biofilm formation and swarming motility

Persistence is the ability of a bacterial population (or a part of it) to survive exposure to bactericidal drugs^46^, without becoming resistant to the antibiotic. Toxin-antitoxin systems have been described to play a role in triggering persistence in bacteria by reversibly intoxicating the bacterium and producing a transient arrest of metabolic activity that renders antibiotics ineffective. This role is controversial, at least for some TAs, since several studies were later shown to be biased by the activity of prophages^47^. Given the deletion of a variety of TS systems in the ΔSI mutant we decided to test persistence upon exposure to high concentrations of antibiotics. We challenged growing cultures of *V. cholerae* WT and ΔSI to 10-fold the minimum inhibitory concentration (MIC) of ampicillin and ciprofloxacin, and viability was measured along a 6 h-period (Figure 8A). Similar decreases in the number of viable cells were observed for both strains, of varying intensities depending on the antibiotic molecule, suggesting that toxin-antitoxin systems encoded in the superintegron do not play a role in antibiotic persistence in *V. cholerae*.

**Figure 8.**
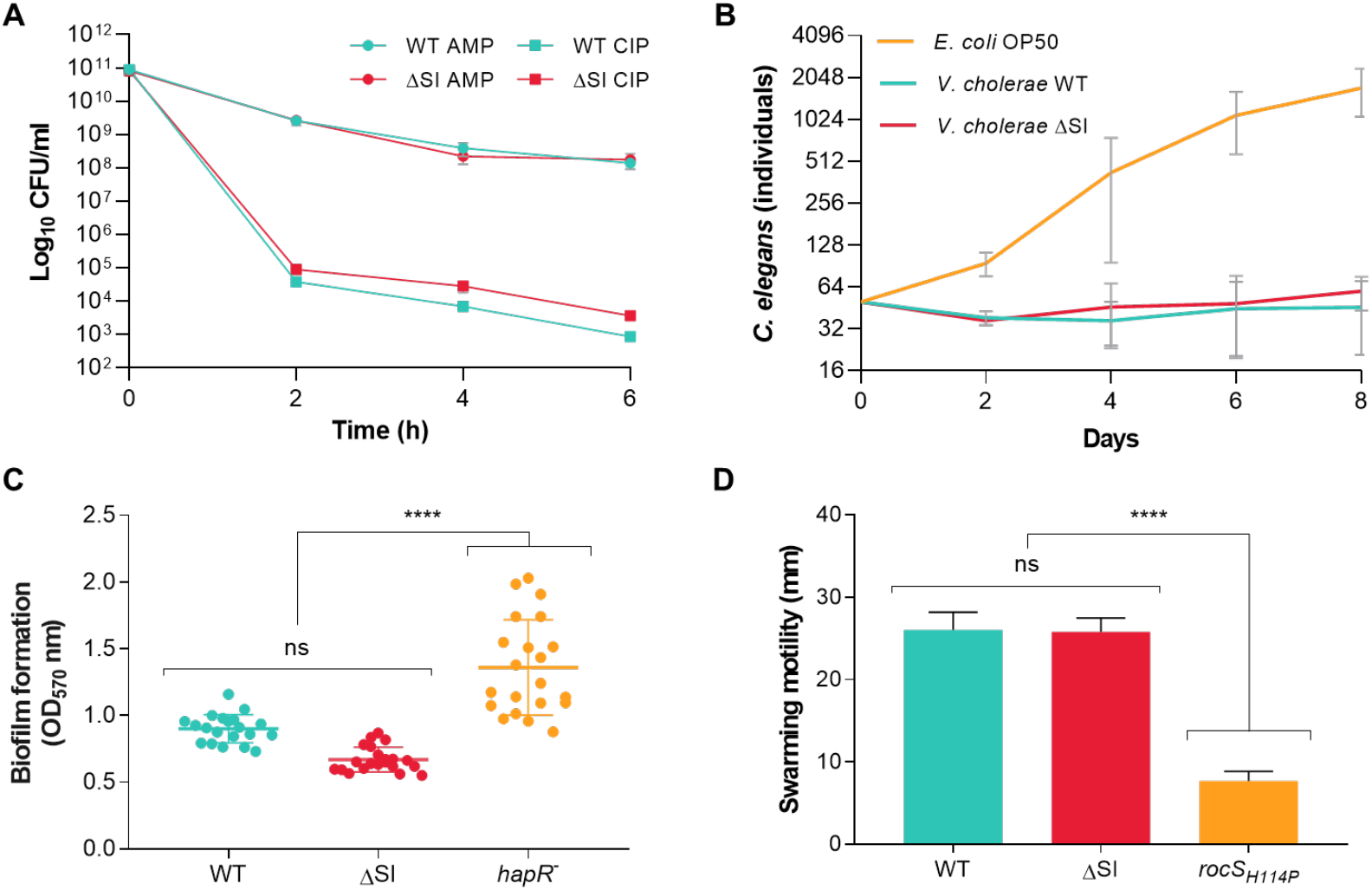
(**A**) Persistence assay of *V. cholerae* WT and ΔSI mutant challenged with 10-fold the MIC of ampicillin and ciprofloxacin, individually. CFUs/ml are plotted against time. Error bars indicate standard deviations from three independent biological replicates. (**B**) Virulence assay of *V. cholerae* WT and ΔSI mutant in a *C. elegans* model. *E. coli* OP50 was used as a negative control. Error bars indicate standard deviations from three independent biological replicates. (**C**) Biofilm formation of *V. cholerae* WT and ΔSI mutant. A *V. cholerae hapR* (frameshift) mutant was included as a positive control of biofilm formation. Data from at least 20 independent colonies is represented. (**D**) Swarming motility assay of *V. cholerae* WT and ΔSI mutant. A *V. cholerae rocS* (H114P) mutant was included as a non-motile control strain. Error bars indicate standard deviation of at least three biological replicates. P values (****) were calculated by pairwise comparison using the one-way ANOVA test (p < 0,0001). Ns: not significant.

*V. cholerae* expresses well-known virulence factors to colonize and cause infection in the mammalian host, as the cholera toxin or the toxin-coregulated pili^48,49^. The nematode *Caenorhabditis elegans* has been widely used as an invertebrate infection model to screen and identify virulence factors of several human pathogens, including *V. cholerae*^50,51^. Toxin-antitoxin systems in the superintegron are reported to be associated with virulence. For instance, the RelBE TA systems of the superintegron play a role in intestine colonization in a mouse model^52^. However, evidence on the implication of other elements in the superintegron in the virulence of *V. cholerae* is lacking. To address this question, we performed a *C. elegans* killing assay. Hermaphrodites L4 worms raised on *Escherichia coli* OP50 were transferred onto bacterial lawns of WT and ΔSI strains. The number of *C. elegans* individuals were counted every 48 h for 8 days. *E. coli* OP50 was used a control. As shown in Figure 8B, we did not observe any differences in lethality between *V. cholera*e WT and the ΔSI mutant.

Biofilm formation and motility are key behaviors in the lifestyle of *V. cholerae*, with strong implications in its survival in the environment, pathogenesis, dispersal, and natural competence. Motility is involved in regulating biofilm formation by participating in important processes, as surface attachment or biofilm dispersal^53^. We investigated the role of the SI by measuring biofilm formation and swarming motility. As shown in Figure 8C and 8D, neither process was significantly affected in the ΔSI strain. As controls, we included a *V. cholerae* strain with frameshift mutation in *hapR* (a regulator of quorum sensing) leading to an increase in biofilm production, and a strain with an amino acid change mutation in *rocS* as a non-motile control.

## DISCUSSION

The working model of integrons is based on semiconservative recombination and the strong expression of cassettes upon integration, highlighting that cassettes must be adaptive to be maintained in the array. This, together with the intimate intertwining between integrons and their hosts (through the SOS response controlling the expression of the integrase^54^, or through host factors^29,55^) put integrons at the service of the host to provide adaptation on demand^2,55,56^. A clear example of this is their role in the rise and spread of multidrug resistance during the last decades. In Vibrionaceae, integrons provide genetic diversity upon which natural selection can act^57^. It is therefore assumed that sedentary chromosomal integrons provide great adaptability too. Yet a better understanding of how they do so, remains out of reach because the functions encoded in the cassettes of chromosomal integrons are mostly unknown^15^. Apart from antibiotic resistance genes and toxin-antitoxin systems, only a handful of cassettes have been characterized (for a review see^33^ and^58^).

Chromosomal integrons have co-evolved with their hosts during aeons. It is therefore plausible that hosts have co-evolved with integrons and have somehow adapted to the presence of such massive structures. Also, the functions encoded in cassettes could be intimately linked to the host’s physiology, somehow interfering or modulating housekeeping functions. Indeed, *in silico* analysis of cassettes in Vibrionaceae revealed potential functions like information storage, cellular processes, and metabolism^15^. This is certainly at odds with the variable nature of the array of cassettes, that would imply that cassettes provide exclusively “self-contained” or independent functions, that do not interfere with those encoded in the rest of the genome. Here we have tested the hypothesis that SCIs are genetically and functionally independent of the host by relocating and deleting the superintegron of *V. cholerae* and testing a broad variety of phenotypes of importance in the lifestyle of this bacterium. The result is surprisingly clear: the ΔSI mutant behaves almost identically to the WT strain in most conditions tested, and differences (like those observed in the use of di-peptides) are, at best, very mild. This tilts the balance towards the “plug-and-play” model of cassette function.

Focusing on the subtle differences that we have observed, we note that the deletion of the SI has a limited impact on gene expression, with only three genes (*aspA*, *dcuA* and *rimO*) being mildly affected. *aspA* and *dcuA* are involved in nitrogen metabolism. AspA carries out the reversible conversion of L-aspartate to fumarate, releasing ammonia for nitrogen assimilation^59^. This enzyme has a specific interaction with DcuA, which functions as a L-aspartate/fumarate antiporter under both aerobic and anaerobic conditions^60^. This interaction has been proposed to form a metabolon, where DcuA facilitates the uptake of aspartate, while AspA converts it to fumarate^61^. The GSEA analysis revealed an enrichment in arginine and isoleucine biosynthetic processes in the ΔSI strain. Arginine synthesis is derived from glutamate, which, in turn, originates from L-aspartate. Additionally, the precursor molecule for isoleucine biosynthesis is pyruvate, a product of glycolysis, and an indirect byproduct of the aspartate metabolism ^62^. These findings, coupled with the observation that the ΔSI mutant exhibits an impaired growth when utilizing certain dipeptides and tripeptides as nitrogen sources, may suggest a potential connection between these processes. However, it is important to note that these pathways are not directly linked, and further experimental data would be needed to ascertain whether a change in metabolic balance has occurred within the cell. Given the subtle signal from our data, we remain cautious about the biological significance of these observations.

The obtention of a ΔSI mutant of *V. cholerae* is a long-awaited milestone in the field, unblocking the study of sedentary chromosomal integrons. Getting rid of the interference of the cassettes within the SI allows to use this strain as a chassis to easily perform many of the experiments that have been delivered in *E. coli* with the class 1 integron, allowing to enrich the field with data from other integron models. It is also the perfect chassis to unveil cassette functions in a classical *in trans* approach, indeed one of the key questions that remains unanswered in the field and one with an important biotechnological potential.

## DATA AVAILABILITY

The RNA-seq data included in this publication have been deposited in NCBI’s Gene Expression Omnibus^63^ and are accessible through GEO Series accession number GSE247496.

## SUPPLEMENTARY DATA

Supplementary Data are available online at the publisher’s web.

## AUTHOR CONTRIBUTIONS

Paula Blanco: conceptualization, formal analysis, methodology, validation, writing-original draft. Filipa Trigo: methodology, analysis and validation. Laura Toribio-Celestino: methodology, analysis, validation. Lucía García-Pastor: methodology, analysis and validation. Niccolò Caselli: methodology, analysis and validation. Francisco Ojeda: methodology, analysis and validation. Baptiste Darracq: methodology, analysis and validation. Ole Skovgaard: methodology, analysis and validation. Álvaro San Millán: formal analysis, writing-review and editing. Didier Mazel: conceptualization, writing-review and editing. Céline Loot: conceptualization, analysis, validation, writing-review and editing. José Antonio Escudero: conceptualization, methodology, formal analysis, validation, writing-original draft, review and editing.

## ACKNOWLEDGEMENTS

Authors would like to thank Laurence Van Melderen, Nathalie Balaban and David Bikard for helpful discussion, and Claire Vit for technical assistance.

## FUNDING ENTITIES

This work is supported by the European Research Council (ERC) through a Starting Grant [ERC grant no. 803375-KRYPTONINT;]; Ministerio de Ciencia, Innovación y Universidades [BIO2017-85056-P, PID2020-117499RB-100]; JAE is supported by the Atracción de Talento Program of the Comunidad de Madrid [2016-T1/BIO-1105 and 2020-5A/BIO-19726]; PB is supported by the Juan de la Cierva program [FJC 2020-043017-I]; FTR is supported by the Portuguese Fundação para Ciência e a Tecnologia [SFRH/BD/144108/2019]; LTC and ASM are supported by the European Research Council (ERC) under the European Union’s Horizon 2020 research and innovation program (ERC grant no. 757440-PLASREVOLUTION). BD, DM and CL are supported by Institut Pasteur, the Centre Nationale de la Recherche Scientifique (CNRS-UMR3525) and the Fondation pour la Recherche Médicale (FRM Grant No. EQU202103012569)

## CONFLICTS OF INTEREST

Universidad Complutense de Madrid and Institut Pasteur have filed a patent covering SeqDelTA and the *V. cholerae* ΔSI strain B522. Patent number: P202330947. Inventors: PB, FTR, DM, JAE.

## MATERIALS AND METHODS

### SI relocation from Chr2 to Chr1

All strains, plasmids and primers used for the SI relocation are listed in Supplementary Table S1, S4, and S5. To relocate the whole superintegron, we used a genetic tool developed in the lab based on the recombination of two bacteriophage attachment sites^64^. Prophage excision from the host chromosome relies on site-specific recombination between two sequences flanking the phage, termed attachment sites *attL* and *attR*. This recombination is carried out by phage-specific recombinases “Int” and their cognate RDF (recombination directionality factor), or excisionase, Xis. The tool developed in the lab uses the attachment sites from bacteriophages HK and λ (Supplementary Figure 2). The *attL/attR* pairs of each site were associated with parts of a genetic marker that is reconstituted when the sites recombine: HK sites were associated with parts of a *bla* gene (ampicillin resistance) and the λ sites with parts of *lacZ*. Co-expression of the HK and λ integrases/excisionases triggers the recombination of the partner *attL/attR* sites, which allows for the reconstitution of fully functional *bla* and *lacZ* markers and the selection of clones in which the SCI relocation has occurred. Blue colonies growing in LB supplemented with carbenicillin and X-gal (5-bromo-4-chloro-3-indolyl-β-D-galactopyranoside) were PCR verified and used for growth curves.

### SeqDelTA

All strains, plasmids, and primers for the deletion of the SI are listed in Supplementary Table S2, S3, S4, and S5. To generate the ΔSI mutant of *V. cholerae* N16961, we developed SeqDelTA, as depicted in Figure 2. After identifying the 19 toxin-antitoxin (TA) systems contained in the *V. cholerae* N16961 superintegron, we prepared successive constructs designed with homology regions and a selective marker. A left homology region (LHR) was designed and maintained throughout the procedure. This region corresponded to the *catB9* gene (VCA0300), and was amplified with primers VCA 299 F and LRHI VCA 300 R. For the successive deletions, different right homology regions (RHD) were designed to maintain antitoxin gene but to eliminate part of the corresponding toxin gene, ensuring that the event was not lethal to the bacteria (Figure 2 and Table S2). As selective markers, three different antibiotic resistance genes (Zeo^R^, Cm^R^, and Carb^R^) were used and amplified with the same primers, facilitating the consequent design and assembly of the HRs. These three fragments (LHR, resistance marker, and RHR) were then assembled by SOE-PCR, placing the resistance marker between the LHR and the RHR. Each successive construct was introduced by natural transformation (described below) into the *Vibrio cholerae* strain obtained from the previous TA system deletion. This strategy allowed the removal of the resistance marker introduced in the previous step while advancing the deletion of TA systems by changes in the RHR. When necessary, due to gene orientation and/or gene order in the TA system, alternative LHRs were designed, and two resistance markers were maintained in consecutive allelic replacements before the new step allowed deletion of both (Table S2).

The last deletion was performed using the R6K origin of replication-plasmid pMP7 as described in^34,65^. pMP7 replication is dependent of the cell π protein and encodes the *ccdB* toxin under the control of the inducible promoter P_BAD_. Five hundred bp of each side of the superintegron were amplified with primers RHI link pMP7 F and LRHI+D R for the LHR, and RHD F and RHD link pMP7 R for the RHR. The two fragments were then joined by SOE-PCR and digested with enzymes NaeI y HindIII. Finally, the insert was ligated with the empty pMP7 backbone, giving rise to pMP7_Δ*intIA attIA zeo^R^*. We started the cloning steps by transforming the pMP7_Δ*intIA attIA zeo^R^* plasmid into *ccdB*-resistant *E. coli* π3813 (*thy::pyr* +) competent cells. Transformants were selected in LB agar plates containing Cm (25 μg/ml), thymidine (dT; 0,3 mM) and glucose (1%). For conjugal transfer of pMP7_Δ*intIA attIA zeo^R^* into *V. cholerae*, the plasmid was first transformed into the donor strain *E. coli* β3914 (*dap::pir* +). Transformants were selected in LB agar plates supplemented with Cm (25 μg/ml), 2,6-diaminopimelic acid (DAP; 0,3 mM) and glucose (1%). Conjugation assay was performed by growing both donor (*E. coli* β3914) and recipient (*V. cholerae* A066) strains to a OD_600_ of 0,2. Then, cells were mixed in a ratio of 1/10 (donor/recipient) and transferred to a mating filter placed on a LB agar plate supplemented with DAP (0,3 mM) and glucose (1%) and conjugation was performed overnight at 37°C. Integration of the pMP7_Δ*intIA attIA zeo^R^* into the *Vibrio* genome was selected by washing the filter in 5 ml of LB and plating serial dilutions on Cm (2,5 μg/ml) and glucose (1%) plates, but lacking DAP. Afterwards, Cm-resistant colonies were grown in liquid LB medium and plated in LB agar plates supplemented with L-arabinose (0,2%) to express *ccdB* and select for the second crossover that implies the excision of the pMP7 backbone. After this process, we Sanger-sequenced the superintegron region and selected a *V. cholerae* ΔSI strain (A101).

### Natural transformation assays

Transformation assays were performed as previously described in^66^ with some modifications. Briefly, strains were inoculated from frozen stocks into LB broth and grown rolling overnight at 30°C. Overnight cultures were diluted 1:100 and grown to an OD_600_ of 1.0. Cells were then washed and resuspended in Instant Ocean^®^ 0,5x (IO), and inoculated in a final volume of 1mL IO with 150 μl of chitin (Apollo Scientific^®^) slurry. Cells were incubated in static conditions at 30°C for ∼18 hours. Then, 550 μl of supernatant was removed from the reactions, and the purified transforming DNA (tDNA) was added. DNA and cells were incubated for ∼18 hours in static conditions at 30°C. When needed, reactions were outgrown by adding 1 m LB broth and shaking at 30°C for 2 hrs. Cells were plated in LB agar with the corresponding antibiotic.

### DNA isolation, WGS, and data analysis

Genomic DNA of the obtained *V. cholerae* ΔSI strain (A101) was extracted using the DNeasy® Blood and Tissue Kit (QIAGEN) following the manufacturer’s protocol. The DNA quantity was determined using a BioSpectrometer (Eppendorf). DNA sequencing was performed at Institut Pasteur using Illumina paired-end reads, at MiGS (Pittsburgh, US), now SeqCoast Genomics (Portsmouth, US) and inhouse at MBA (Madrid) using MinIon. Data analysis was accomplished with Geneious Prime software and the genetic variants were identified by mapping the generated reads to the *V. cholerae* N16961 reference genome (Accession number: chromosome 1 CP028827.1; chromosome 2 CP028828.1).

### Marker Frequency Analysis (MFA)

Marker Frequency Analysis was performed as described in^35^. Cells were grown in LB at 37°C with shaking and genomic DNA was isolated using the DNeasy® Tissue Kit (Qiagen) from exponential and stationary phase cultures. Genomic DNA was sequenced with Illumina technology. Short reads were aligned to the reference genome using Bowtie2^67^ and R2R. The number of reads starting per bp (N) were calculated for 1 kbp and 10 kbp windows. The exponential phase data were normalized with a factor calculated as the deviation of observed N from calculated N for each window of the stationary phase culture data. Any window including repeated sequences was excluded and N was plotted as a function of the absolute position, centered on the replication origins.

### Correction of *V. cholerae* ΔSI mutations

All the strains, plasmids and primers are listed in Supplementary Table S3, S4 and S5. Three mutations were found in *V. cholerae* ΔSI strain (A101): *rocS* (VC0653), *cry2* (VC01392) and *rpoS* (VC0534). While *rocS* and *rpoS* mutations led to frameshifts, *cry2* contained a non-synonymous mutation. To revert the mutations to the WT variant, allelic exchanges were performed using the pMP7, as previously described. To provide homology for the allelic replacement, 1.000 bp-fragments of the N16961 strain genes *rocS*, *rpoS* and *cry2* were amplified with primers *rocS*_pMP7 F/R, *rpoS*_pMP7 F/R, and *cry2*_pMP7 F/R, respectively, which contain 20 bp-homology fragments with the pMP7 cloning site; pMP7 backbone was amplified with oligos pMP7_bb_Gibson F/R. Cloning of the amplified fragments was performed using Gibson assembly^68^, giving rise to pMP7_*rocS*, pMP7_*rpoS*, and pMP7_*cry2*. We performed the mating and allelic exchanges as previously mentioned, starting with pMP7_*rocS*, then pMP7_*rpoS*, and finally, pMP7_*cry2*. After each step, we Sanger-sequenced the targeted gene and selected a *V. cholerae* strain corrected for *rocS*+ (A677), *rocS*+, *rpoS*+ (A684), and finally *rocS*+, *rpoS*+, *cry2*+ (B522). We verified by Illumina whole-genome sequencing that *V. cholerae* ΔSI (B522) did not contain any other unintended mutations.

### RNA extraction and preparation for RNA-seq

Total RNA was extracted from *V. cholerae* WT and ΔSI cultures grown in LB at 37°C in both exponential (OD_600_ 0,8) and stationary (OD_600_ 2,8) growth phases using the RNeasy® Mini Kit (QIAGEN), following the manufacturer’s protocol. To eliminate any residual DNA, RNA was treated using the TURBO DNA-free™ Kit (Invitrogen). RNA concentration was measured using a BioSpectrometer (Eppendorf), and RNA integrity was determined using a Qubit™ 4 fluorometer (Invitrogen) with the RNA IQ Assay Kit (Invitrogen). Ribodepletion RNA library and sequencing was performed at the Oxford Genomics Centre using a NovaSeq6000 sequencing system (Illumina). Three biological samples per condition were sequenced.

### RNA-seq data analysis

Raw Illumina reads were trimmed and adapter-removed using Trim Galore v0.6.6 (https://github.com/FelixKrueger/TrimGalore) with a quality threshold of 20 and removing reads shorter than 50 bp. Trimmed paired reads were mapped to the *V. cholerae* N16961 reference genome (Accession number: chromosome 1 CP028827.1; chromosome 2 CP028828.1) using BWA-MEM v0.7.17 ^69^. featureCounts from the Rsubread v2.10.2 package ^70^ was used to obtain read counts per annotated feature, including CDS, ncRNA, tmRNA, RNase P, tRNA, antisense RNA and SRP RNA. Differential expression analysis was performed from raw counts using DESeq2 v1.36.0^71^, by comparing the expression data of the deletion mutant against the WT strain.

Differentially expressed genes (DEGs) of CDSs were annotated with Gene Ontology (GO) terms retrieved from UniProt on July 7, 2023^72^. Gene Set Enrichment Analysis was performed to identify sets of biological processes, molecular functions, or cellular components with different expression profiles between the superintegron deletion mutant and WT strains, in both exponential and stationary phases. For this, a pre-ranked list of DEGs, ordered by log2 fold changes, was provided to the GSEA function of the clusterProfiler v4.8.1^73^. Genes belonging to the superintegron (positions 312,057-438,942 bp of chromosome 2) were not included to prevent biased enrichment results caused by the acute downregulation of the superintegron genes. GSEA was run with default parameters, correcting for multiple tests with the Benjamini-Hochberg procedure.

### Quantitative real-time PCR

Independent samples of total RNA from *V. cholerae* WT and ΔSI cultures grown in LB at 37°C in both exponential (OD_600_ 0,8) and stationary (OD_600_ 2,8) were isolated as described previously. To eliminate any residual DNA, RNA was treated using the TURBO DNA-free™ Kit (Invitrogen) and RNA concentration was measured using a BioSpectrometer (Eppendorf). Before cDNA synthesis, a previous DNA wipe-out step was performed using the QuantiTect® Reverse Transcription Kit (QIAGEN) following the manufacturer’s instructions; cDNA synthesis was carried out afterwards using the same kit using the temperature steps: 42°C for 15 min; 95 °C for 3 min.

RT-qPCR was performed in an Applied Biosystems QuantStudio 3 using the Fast SYBR™ Green Master Mix (Thermo Fischer Scientific). Primers used are listed in Supplementary Table S5, and *gyrA* was used as housekeeping gene. Relative changes in gene expression for the superintegron deletion mutant with respect to the WT were determined according to the threshold cycle method (2−^ΔΔCT^). Mean values were obtained from two independent biological replicates with three technical replicates each.

### Note on the controls used for the search for phenotypes

The original *V. cholerae* N16961 isolate contains mutations in the *hapR* gene and is not naturally competent. The deletion of the superintegron has been performed in a well-known derivative strain in which natural competence is restored by the insertion of a transposon carrying a functional *hapR* gene (strain A001 in the Supplementary Table S3). It is known that *V. cholerae* cultures have a tendency towards the *hapR*^-^ genotype. Through genome sequencing we observed this phenomenon in our stock of the parental strain, which seemed to rise through homologous recombination between both *hapR* alleles. Since *hapR*^-^ strains have certain phenotypes, such as distinct growth curves or a decreased virulence, we performed most phenotypic experiments using as a control a strain of *V. cholerae* N16961, in which mutations in *hapR* have been corrected without the addition of a second allele (A096 in our strain list). In RNAseq and MFA experiments were we could use as a control the parental strain (A001), by confirming through sequence analysis that all replicates contained a functional *hapR* allele.

### Growth curves

Overnight cultures were prepared by inoculating single colonies from LB agar plates in LB liquid medium and incubating for 16-20 h at 37°C. Cultures were then diluted 1:1.000 in fresh medium and 200 μl were transferred into 96-well plates (Nunc, Thermo Scientific). OD_600_ was measured every 15 min for 16 h using a Biotek Synergy HTX plate reader (Agilent). Vmax and area under the curve (AUC) were determined using Gen5 software and MATLAB, version R2022a (Mathworks), respectively. At least 24 independent replicates were included in the experiment and significant differences were assessed by performing unpaired t-test.

Growth curves in the presence of subinhibitory concentration of norfloxacin, ceftriaxone and tannic acid were performed by diluting the compounds at the desired concentrations in MH medium. Overnight cultures of *V. cholerae* WT and ΔSI were diluted 1:1000 in the antimicrobial-containing MH media in 96-well plates (Nunc, Thermo Scientific). OD_600_ was measured every 20 min for 24 h using a Biotek Synergy HTX plate reader (Agilent). At least three biological replicates were included in the experiment.

### Competition assays

Competition assays were performed by flow cytometry to measure the fitness values of *V. cholerae* WT and ΔSI mutant relative to an *E. coli* DH5α strain as a common competitor carrying a pSU38 plasmid with the *gfp* under the control of the constitutive promoter P_C_S. Competition procedure was performed as described in^74^ with some modifications. Briefly, pre-cultures were prepared by inoculating single colonies from a LB agar plate in 200 μl of liquid LB in a 96-well plate (Nunc, Thermo Scientific). After 22 h of growth at 37°C with 250 rpm shaking, cultures were mixed at 1:1 proportion and diluted 1:400 in fresh medium. In order to confirm the initial proportions of *E. coli* carrying *gfp* and the non-fluorescent *V. cholerae* strains, cells were diluted 1:400 in NaCl (0,9%) and 30.000 events per sample were recorded using a CytoFLEX S flow cytometer (Beckman Coulter). Bacterial mixtures were competed for 22 h at 37°C with 250 rpm shaking, and final proportions were determined as stated above. The fitness values of both *V. cholerae* WT and ΔSI, relative to *E. coli* P_C_S::*gfp* were calculated using the formula: w=ln(N_final,gfp-_/N_initial,gfp-_)/ln(N_final,gfp+_/N_initial,gfp+_), where w is the relative fitness of the non GFP-tagged *V. cholerae* strains, N_initial,gfp-_ and N_final,gfp-_ are the numbers of non GFP-tagged *V. cholerae* at the beginning and end of the competition, and N_inital,gfp+_ and N_final,gfp+_ are the numbers of *E. coli* P_C_S::*gfp* cells at the beginning and end of the competition, respectively. Twelve biological replicates were performed for each competition experiment and significant differences were assessed by performing unpaired t-test.

### Phenotypic microarray assay

Phenotype Microarrays (Biolog) PM1-PM20 were tested as previously described in^45^ with some modifications. Briefly, single colonies of *V. cholerae* WT and ΔSI were inoculated in liquid LB medium and grown overnight at 37 °C. For PM1 and PM2 plates inoculation, 120 μl of Redox Dye mix D (100x), 1 ml of sterile water and 880 μl of a 85% T cell suspension in NaCl (0.9%) were added to 10 ml of IF-0 (1.2x), resulting in a final volume of 12 ml per plate. For PM3-PM8 plates inoculation, which requires an appropriate carbon source, 120 μl of sodium pyruvate (2M) was added as an additive and mixed with 120 μl of Redox Dye mix D (100x), 880 μl of sterile water, and 880 μl of a 85% T cell suspension in NaCl (0.9%). Then, this solution was added to 10 ml of IF-0 (1.2x), resulting in a final volume of 12 ml per plate. For PM9-PM20 plates inoculation, 120 μl of Redox Dye mix D (100x), 1 ml of sterile water and 880 μl of 1:200 dilution of the 85% T cell suspension in NaCl (0.9%) were added to 10 ml of IF-10 (1.2x), resulting in a final volume of 12 ml per plate. Each plate was inoculated with 100 μl per well of the respective prepared solution and then sealed using Breathe-Easy® sealing membranes (Sigma-Aldrich). Plates were then transferred to a Biostack Microplate Stacker (Agilent) to process several plates per day and incubated at 37°C using a Memmert IF750 (Memmert). OD_590_, which indicates the reduction of the tetrazolium dye, was measured every 20 min during 24 h using a Biotek Synergy HTX plate reader (Agilent). Experiments were conducted in duplicate and growth characteristics were determined by calculating the AUC using MATLAB, version R2022a (Mathworks).

### Persistence assay

Three colonies of *V. cholerae* WT and ΔSI were inoculated in LB liquid medium and grown overnight at 37°C. Overnight cultures were diluted 1:100 in 1 ml of fresh LB medium in 24-well plates and grown at 37°C on 150 rpm shaker for 3h. After 3 h incubation, cultures were challenged individually with 10-fold MIC of ciprofloxacin (MIC = 0,008 μg/ml) or ampicillin (MIC = 16 μg/ml) and incubated for 6 h at 37°C on 150 rpm shaker. The CFUs/ml were enumerated at times 0 h (before adding the antibiotics), 2 h, 4 h, and 6 h by serially diluting the cells and plating on LB agar.

### Caenorhabditis elegans virulence assay

*C. elegans* N2 Bristol individuals were routinely maintained at 20°C on potato dextrose agar (PDA; Sigma-Aldrich) plates seeded with *E. coli* OP50. Overnight LB cultures of the *V. cholerae* strains and *E. coli* OP50 were diluted 1:100 and grown at 37°C until an OD_600_ of 0,8 was reached. Then, 50 μl from each strain were spread on 6-cm-diameter plates containing PDA medium. Plates were incubated for 16-20 h at 37°C to form a bacterial lawn and 50 L4-stage hermaphrodite individuals were then placed on the PDA plates and incubated at 20°C. Live *C. elegans* were scored every 48 h for 8 days. An individual was considered dead when it no longer responded to touch. *E. coli* OP50 was used as a negative control of virulence. At least, three independent experiments were performed.

### Biofilm formation assay

Overnight cultures were diluted 1:100 in fresh LB medium. Hundred microliters were transferred to into 96-well Serocluster™ plates (Costar) and incubated at 37°C for 20 h without shaking. After incubation, all the wells were washed with sterile distilled water for three times and dried. Biofilm mass was stained with 125 μl cristal violet (1%) per well and incubated for 15 min at room temperature. Cristal violet was removed by performing three washes with sterile distilled water. The stained biofilm mass was then detached from the wells using acetic acid (30 %) and transferred to a new 96-well plate (Nunc, Thermo Scientific). OD_570_ was measured using a Biotek Synergy HTX plate reader (Agilent). At least 20 independent experiments were performed per strain. One way-ANOVA was used to determine significant differences among the three groups.

### Swarming motility assay

Swarming motility assays were performed as described in^75^. Briefly, 3 μl of overnight cultures were soaked on motility agar plates containing 10 g/l tryptone, 5 g/l NaCl, and 0,25% bacto-agar. The swarming motility halos were measured and compared after growth at 37°C for 6 h. At least three independent experiments were performed per strain. One way-ANOVA was used to determine significant differences among the three groups.

