## Supplementary Material for "Chromosomal Integrons are Genetically and Functionally Isolated Units of Genomes"

**Table S1.** Strains generated for the relocation of the superintegron

| Name | Genotype | Reference | Lab collection |
|---|---|---|---|
| **8637** | *V. cholerae* N16961 biovar El Tor, *hapR*+ (Tn::*hapR*) [StrepR] WT | Lab Collection from [1] | PGB |
| **B805** | 8637 VC0018-*bla*3'-attL$_{HK}$-*cat*-attR$_{\lambda}$ *lacZ*-3'-VC0019 | J. Bland, unpublished | PGB |
| **B825** | 8637 VC0019-*bla*3'-attL$_{HK}$-*cat*-attR$_{\lambda}$ *lacZ*3' -VC0018 | J. Bland, unpublished | PGB |
| **D282** | 8637 rplT-attR$_{HK}$-*bla*5'-*aph* | J. Garriss, unpublished | PGB |
| **F854** | B805 VC0018-*bla*3'-attL$_{HK}$-*aadA7*-attR$_{\lambda}$ *lacZ*3'-VC0019 | This study, replacement by natural transformation of the *cat* fragment of B805 by the *spec* fragment from pF849 | PGB |
| **F855** | B825 VC0019-*bla*3'-attL$_{HK}$-*aadA7*-attR$_{\lambda}$ *lacZ*3' -VC0018 | This study, replacement by natural transformation of the *cat* fragment of B825 by the *spec* fragment from pF849 | PGB |
| **F875** | F854 rplT-attR$_{HK}$-*bla*5'-*aph*-*intIA* | This study, insertion by natural transformation of the *aph* fragment from pF851 | PGB |
| **F876** | F855 rplT-attR$_{HK}$-*bla*5'-*aph*-*intIA* | This study, insertion by natural transformation of the *aph* fragment from pF851 | PGB |
| **J060, J061, J183** | F875 VCA0508-*ble*-*lacZ*5'-attL$_{\lambda}$-VCA0510 | This study, insertion by natural transformation of the *ble* fragment from pF912 | PGB |
| **J062, J063, J184** | F876 VCA0508-*ble*-*lacZ*5'-attL$_{\lambda}$-VCA0510 | This study, insertion by natural transformation of the *ble* fragment from pF912 | PGB |
| **J325, J327, J333** | J060, J061 and J183 with pF930 | This study, transformation of J060, J061 and J183 by pF930 | PGB |
| **J329, J331, J335** | J062, J063 and J184 with pF930 | This study, transformation of J062, J063 and J184 by pF930 | PGB |
| **J395, J399, J407 (SCI lag)** | SCI relocated near ori1 (Chr1) betweenVC0018 and VC0019. *attCbs* carried by the lag strand template. From J329, J331, J335 | This study, relocation by site-specific recombination of the SCI. | PGB |
| **J387, J391, J403 (SCI lead)** | SCI relocated near ori1 (Chr1) between VC0018 and VC0019. *attCbs* carried by the lead strand template. From J325, J327, J333 | This study, relocation by site-specific recombination of the SCI. | PGB |

**PGB:** Plasticité du Génome Bactérien Lab.

**Table S2.** Strains generated with SeqDelTA

| Name | Genotype | Primers LHR | Primers RHR | Reference | Lab collection |
|---|---|---|---|---|---|
| **A400** | ΔVCA300-311. [Zeo$^R$] | VCA 299 F – LRHI VCA 300 R | LRHD VCA 311 F – VCA 312 R | This study | MBA |
| **A003** | ΔVCA300-320. [Cm$^R$] | VCA 299 F – LRHI VCA 300 R | LRHD VCA 318 F – VCA 320 R | This study | MBA |
| **A004** | ΔVCA300-325. [Carb$^R$] | VCA 299 F – LRHI VCA 300 R | LRHD VCA 323 F –VCA 325 R | This study | MBA |
| **A005** | ΔVCA300-332. [Zeo$^R$] | VCA 299 F – LRHI VCA 300 R | LRHD + prom VCA 332 F – prom VCA 332 R VCA 333 F – VCA 334 R | This study | MBA |
| **A006** | ΔVCA300-350. [Carb$^R$] | VCA 299 F – LRHI VCA 300 R | LRHD VCA 348 F – VCA 350 R | This study | MBA |
| **A007** | ΔVCA300-360. [Cm$^R$] | VCA 299 F – LRHI VCA 300 R | LRHD VCA 359 F – VCA 360 R | This study | MBA |
| **A009** | ΔVCA300-385. [Zeo$^R$] | VCA 299 F – LRHI VCA 300 R | LRHD VCA 311 F – VCA 386 R | This study | MBA |
| **A023** | ΔVCA300-391. [Carb$^R$] | VCA 299 F – LRHI VCA 300 R | LRHD VCA 391 F –VCA 393 R | This study | MBA |
| **A024** | ΔVCA300-391 + VCA422-444. [Carb$^R$, Zeo$^R$] | VCA 421 F – LRHI VCA 422 R | LRHD VCA 443 F –VCA 447 R | This study | MBA |
| **A029** | ΔVCA300-470. [Cm$^R$] | VCA 299 F – LRHI VCA 300 R | LRHD + prom VCA 469 F – prom VCA 469 R VCA 469 F – VCA 470 R | This study | MBA |
| **A035** | ΔVCA300-470 + VCA474-483. [Cm$^R$, Zeo$^R$] | VCA 474 F – link AT/TA R / link AT/TA F – LRHI VCA 478 R | LRHD VCA 318 F –VCA 483 R | This study | MBA |
| **A041** | ΔVCA300-483. [Carb$^R$] | VCA 299 F – LRHI VCA 300 R | LRHD VCA 318 F –VCA 483 R | This study | MBA |
| **A047** | ΔVCA300-483 + ΔVCA495- | VCA 494 F – LRHI VCA 495 R | LRHD + prom VCA 497 F – prom VCA 497 R | This study | MBA |

| | | | | | |
|---|---|---|---|---|---|
| | 497. [Carb$^{R}$, Cm$^{R}$] | | VCA 498 F – VCA 499-500 R | | |
| **A054** | ΔVCA300-483 + ΔVCA487-505. [Carb$^{R}$, Zeo$^{R}$] | VCA 486 F – LRHI VCA 487 R | LRHD VCA 348 F –VCA 505 R | This study | MBA |
| **A066** | ΔVCA300-VCA505. [Zeo$^{R}$] | VCA 299 F – LRHI VCA 300 R | LRHD VCA 348 F –VCA 505 R | This study | MBA |

**MBA:** Molecular Basis of Adaptation Lab.

**Table S3.** Parental strains and strains generated with pMP7

| Name | Genotype | Reference | Lab Collection |
|---|---|---|---|
| **A118** | *E. coli* π3813, *ccdB*-resistant strain. (B410*gyrA*462zei::Tn10): *lacIQ*, *thi1*, *relA1*, *supE44*, *endA1*, *recA1*, *hsdR17*, *gyrA462*, *zei*298::Tn10, Δ*thyA*::(*erm-pir*116) [Erm$^{R}$] .Cloning of R6K vectors. | [2] | MBA |
| **A116** | *E. coli* β3914, *ccdB*-resistant strain. *gyrA462 zei298*::Tn10 [Erm$^{R}$, Km$^{R}$, Tc$^{R}$]. Conjugation of R6K vectors. | [2] | MBA |
| **A097** | *E. coli* β3914 pMP7_Δ *intIA attIA zeo$^{R}$* | This study | MBA |
| **A631** | *E. coli* π3813 pMP7-*rocS*$^{+}$ | This study | MBA |
| **A632** | *E. coli* π3813 pMP7-*rpoS*$^{+}$ | This study | MBA |
| **A951** | *E. coli* π3813 pMP7-*cry2*$^{+}$ | This study | MBA |
| **A633** | *E. coli* β3914 pMP7-*rpoS*$^{+}$ | This study | MBA |
| **A634** | *E. coli* β3914 pMP7-*rocS*$^{+}$ | This study | MBA |
| **A982** | *E. coli* β3914 pMP7-*cry2*$^{+}$ | This study | MBA |
| **A001** | *V. cholerae* N16961 biovar El Tor, *hapR*+ (Tn::*hapR*) [Strep$^{R}$]. WT (same as 8637, but stock from MBA lab). | Lab Collection (from [1]) | MBA |
| **A096** | *V. cholerae* N16961 biovar El Tor *hapR*+ (allele repaired *in situ*). WT | [3] | MBA |
| **A101** | *V. cholerae* N16961 ΔSI | This study | MBA |
| **A677** | *V. cholerae* N16961 ΔSI *rocS*$^{+}$ | This study | MBA |
| **A684** | *V. cholerae* N16961 ΔSI *rocS*$^{+}$ *rpoS*$^{+}$; pMP7_*rpoS*$^{+}$ | This study | MBA |
| **B522** | ΔSI *V. cholerae* N16961 ΔSI *rocS*$^{+}$ *rpoS*$^{+}$ *cry2*$^{+}$; pMP7-_*cry2*$^{+}$ | This study | MBA |

**MBA:** Molecular Basis of Adaptation Lab.

**Table S4.** Plasmids

| Name | Plasmid description | Relevant properties and construction | Lab collection |
|---|---|---|---|
| **pD060** | pSW23T::$attC_{aadA7}$ (bs) | $oriV_{R6K\gamma}$, $oriT_{RP4}$; [Cm$^{R}$] [4] | PGB |
| **pA298** | pFLP3 ::*aadA7* | [Sp$^{R}$] Val, unpublished | PGB |
| **pJB6** | pSU38Δ::attR$_{HK}$-attL$_{\lambda}$ | *ori*P15A, [Carb$^{R}$] [5] | PGB |
| **pA401** | pSC101*rep*$^{TS}$:: $int_{HK}$-$xis_{HK}$ | *oripSC101rep*$^{TS}$ $oriT_{RP4}$; [Sp$^{R}$] (Bland, | PGB |

| | | | |
|---|---|---|---|
| | | unpublished) | |
| **pF850** | pTOPO*::VCA0508-ble-bla*3'-*att*LHK-*VCA0510* | *ori*ColE1; [$Km^R$] [6] | PGB |
| **pF912** | pTOPO-VCA0508-*ble-lacZ*5'-$attL_{\lambda}$-VCA0510 | Assembly of three fragments by PCR. Fragments were amplified from pF850 with o4659 and o4708 primers, from pJB6 with o4707 and o4669 and, from 8637 with o4670 and o4662. | PGB |
| **pF851** | pTOPO-rplT-$attR_{HK}$-*bla*5'-*aph-intIA* | Assembly of three fragments by PCR. Fragments were amplified from 8637 with o4286 and o4672 primers, from D282 with o4671 and o4674 and, from 8637 with o4673 and o4302. | PGB |
| **pF849** | pTOPO- VC0018-*bla*3'$attL_{HK}$-*aadA7*-$attR_{\lambda}$ *lacZ*3'-VC0019 | Assembly of three fragments by PCR. Fragments were amplified from B805 with o4665 and o4666, from pA298 with o4663 and o4664 and, from B805 with o4668 and o4667. | PGB |
| **pF930** | pA401 ::$Cm^R$ | *oripSC101rep*$^{TS}$ *oriT*$_{RP4}$; [$Cm^R$] insertion of the XhoI/KpnI digested *cat* fragment from pD060 in pA401. | PGB |
| **pMP7** | Suicide conjugative plasmid used for allelic exchange pSW23T-*araC* $P_{BAD}$-*ccdB* | *oriV*$_{R6K\gamma}$*, oriT*$_{RP4}$*;* [$Cm^R$]. [3] | MBA |
| **pMP7_Δ *intIA attIA zeo*$^R$** | pMP7-derivative used for performing the last deletion step of the superintegron (ΔVCA 291-300) | *oriV*$_{R6K\gamma}$*, oriT*$_{RP4}$*;* [$Cm^R$]. It contains two 500-bp fragments corresponding to each side of the superintegron. Fragments were amplified from *V. cholerae* N16961 gDNA with primers RHI link pMP7 F and LRHI+D R for the LHR, and RHD F and RHD link pMP7 R for the RHR. | MBA |
| **pMP7-*rocS*$^+$** | pMP7-derivative used for the correction of the *rocS* mutation | *oriV*$_{R6K\gamma}$*, oriT*$_{RP4}$*;* [$Cm^R$]. It contains a 1000-bp fragment of the *rocS* (VC0653) gene, amplified from *V. cholerae* N16961 gDNA with primers *rocS*_pMP7 F and *rocS*_pMP7 R. | MBA |
| **pMP7-*rpoS*$^+$** | pMP7-derivative used for the correction of the *rpoS* mutation | *oriV*$_{R6K\gamma}$*, oriT*$_{RP4}$*;* [$Cm^R$]. It contains a 1000-bp fragment of the *rpoS* (VC0534) gene, amplified from *V. cholerae* N16961 gDNA with primers *rpoS*_pMP7 F and *rpoS*_pMP7 R. | MBA |
| **pMP7-*cry2*$^+$** | pMP7-derivative used for the correction of the *cry2* mutation | *oriV*$_{R6K\gamma}$*, oriT*$_{RP4}$*;* [$Cm^R$]. It contains a 1000-bp fragment of the *cry2* (VC01392) gene, amplified from *V. cholerae* N16961 gDNA with primers *cry2*_pMP7 F and *cry2*_pMP7 R. | MBA |

**PGB:** Plasticité du Génome Bactérien Lab; **MBA**: Molecular Basis of Adaptation Lab.

**Table S5.** Primers

| Name | Sequence (5’ → 3’) |
|---|---|
| **Relocation of the superintegron** | |
| **4659** | GAGTAAGTGTCGCTATCGAGGTGCTTCAGTCCTGCTCCTC GGCCACGAAG |
| **4708** | ACTTTTGGCGAAAATGAGACGTTGATATGCAATTGTCGGC ACGTAAGAGG |
| **4707** | CCTCTTACGTGCCGACAATTGCATATCAACGTCTCATTTTC GCCAAAAGT |
| **4669** | CTTAGCCGACAATATCTGCCTATAAAATCAAATAATGATT TTATTTTGAC |
| **4670** | GTCAAAATAAAATCATTATTTGATTTTATAGGCAGATATT GTCGGCTAAG |
| **4662** | AGAAGTGCATGTTTCATCTCCCCCT |
| **4286** | GCTGTAGCACGTATGCTTCC |
| **4672** | ACTACTTAGATAGTATTAGTGACCTGGAATTCGTAATCAT GGTCATAGCT |
| **4671** | AGCTATGACCATGATTACGAATTCCAGGTCACTAATACTA TCTAAGTAGT |
| **4674** | ATTATCCCGTCTTTAGCATGGGTTCCGATCTCGCAGCGGT GGTAAGCGCC |
| **4673** | GGCGCTTACCACCGCTGCGAGATCGGAACCCATGCTAAA GACGGGATAAT |
| **4302** | AAGGTAAGGGGGGTAAAAATCGCAC |
| **4665** | GCCGGAAGGGCCGAGCGCAGAAGTG |
| **4666** | TCTAACAATTCGTTCAAGCCGACGCGACAAATGATTTTAT TTTGACTAAT |
| **4663** | ATTAGTCAAAATAAAATCATTTGTCGCGTCGGCTTGAACG AATTGTTAGA |
| **4664** | GTGACCTGTAACAGAGCATTAGCGCCGAAACCTTGCGCTC GTTCGCCAGC |
| **4668** | GATCACACTCGGGTGATTACGATCG |
| **4667** | GCTGGCGAACGAGCGCAAGGTTTCGGCGCTAATGCTCTGT TACAGGTCAC |
| **SeqDelTA** | |
| **3083** | TCTAGGGCGGCGGATTTGTC |
| **3327** | TGCAATTGTCGGCACGTAAG |
| **VCA 299 F** | GGGCGTTAGAGCTTTATTGG |
| **LRHI VCA 300 R** | GACAAATCCGCCGCCCTAGAGAGCTTTATTTACTCGGACG |
| **LRHD VCA 311 F** | CTTACGTGCCGACAATTGCAGGTAAACGCTCAATCAAAGG |
| **VCA 312 R** | CTTTATTACGACAGCCATCGC |
| **LRHD VCA 318 F** | CTTACGTGCCGACAATTGCATGAAACGGTTCCTATCGTGC |
| **VCA 320 R** | CTGCCAAATCAGCATCAAGC |
| **LRHD VCA 323 F** | CTTACGTGCCGACAATTGCACAATGATGTCTGAAATCGGC |
| **VCA 325 R** | ACCTATGAGCTGACCAATGC |
| **LRHD + prom VCA 332 F** | CTTACGTGCCGACAATTGCAAAATATCACCTAACAAGCGC |
| **prom VCA 332 R** | CTGAGATGATCCTGACAATACCGTGCTTTTCGAG |
| **VCA 333 F** | AAAAGCACGGTATTGTCAGGATCATCTCAGTTCGG |
| **VCA 334 R** | TCATTGACCTCATCAAGACC |
| **LRHD VCA 348 F** | CTTACGTGCCGACAATTGCAATTGAGAGATGCTTCTTCCC |
| **VCA 350 R** | GAAACCTTCAGATAGCCGTC |

| | |
|---|---|
| **LRHD VCA 359 F** | CTTACGTGCCGACAATTGCATCACGTAAATCGGCTTTGGC |
| **VCA 360 R** | TGACCCTGCTCATTTCTTGC |
| **LRHD VCA 311 F** | CTTACGTGCCGACAATTGCAGGTAAACGCTCAATCAAAGG |
| **VCA 386 R** | ATACGATGGACGCCATAAGC |
| **LRHD VCA 391 F** | CTTACGTGCCGACAATTGCAGACTTAAGAATTCCACCGGG |
| **VCA 393 R** | TCAAGGTTAAGTTGGCCACG |
| **VCA 421 F** | GGTGATGCGACTCAAAAAGC |
| **LRHI VCA 422 R** | GACAAATCCGCCGCCCTAGATTGTAACGCTAGAGGTGACC |
| **LRHD VCA 443 F** | CTTACGTGCCGACAATTGCATTGAACTGCTGTTGGAGTGG |
| **VCA 447 R** | ATCGAGGGAAACGCATAACC |
| **LRHD + prom VCA 469 F** | CTTACGTGCCGACAATTGCAGCTCTTATCTGAACTAATTCTTGCC |
| **prom VCA 469 R** | GCCGTAAATGGTTAGCAAATCACTGAGATATTCATCTCGG |
| **VCA 469 F** | CCGAGATGAATATCTCAGTGATTTGCTAACCATTTACGGC |
| **VCA 470 R** | TGGTATAGAAGTCCTGTGCC |
| **VCA 474 F** | GGGCGTTATAACCTAATGGG |
| **link AT/TA R** | GGTAGTTGTTTTTTTGCGACAGCGGCCTTGGGAATATCAACGGCTCC |
| **link AT/TA F** | GGAGCCGTTGATATTCCCAAGGCCGCTGTCGCAAAAAAACAACTACC |
| **LRHI VCA 478 R** | GACAAATCCGCCGCCCTAGAACTCCATTCCTTTTGCGCAG |
| **LRHD VCA 318 F** | CTTACGTGCCGACAATTGCATGAAACGGTTCCTATCGTGC |
| **VCA 483 R** | TTCCTGCCAATACTTGCTACC |
| **VCA 494 F** | TCGATGCTGCTTAACGGTGC |
| **LRHI VCA 495 R** | GACAAATCCGCCGCCCTAGAATCATGGCCATCAACTCTCC |
| **LRHD + prom VCA 497 F** | CTTACGTGCCGACAATTGCACGGGCGTTATATGCTTATAGG |
| **prom VCA 497 R** | GGATTTTCACCATTCACCGCATAGGGCATCAATCTCTGGC |
| **VCA 498 F** | GCCAGAGATTGATGCCCTATGCGGTGAATGGTGAAAATCC |
| **VCA 499-500 R** | TTGTGAGACTAAGCTGACCG |
| **VCA 486 F** | CCTATAATCGCTCAACTGACGG |
| **LRHI VCA 487 R** | GACAAATCCGCCGCCCTAGAACGGATTAGACTGACCTTTCC |
| **VCA 505 R** | CCTACACCGACAATGAAACC |
| **RHI link pMP7 F** | TCTGCGAGGCTGGCCGGCGTCCGTCAGCTGCGTCCAAACG |
| **LRHI+D R** | GCGACACTTACTCAAGCTGTCATGGGTTCTTTGCGAAATC |
| **RHD F** | ACAGCTTGAGTAAGTGTCGC |
| **RHD link pMP7 R** | TCAAGCTTATCGATACCGTCATTTGCTTTATGACTCGCGC |
| **Correction of mutations** | |
| ***rocS*_pMP7 F** | CGATAAGCTTGATATCGAATTCCAGAAAAACCCTTGGTGTGC |
| ***rocS*_pMP7 R** | CAGACAATTGACGGCTCTAGAATCAATCAGTTGAGGATTGC |
| ***rpoS*_pMP7 F** | CGATAAGCTTGATATCGAATTCCGACGCGCAAGCACTTCTTT |
| ***rpoS*_pMP7 R** | CAGACAATTGACGGCTCTAGAGTCAGCAATACCGTAACCA |
| ***cry2*_pMP7 F** | GATAAGCTTGATATCGAATTCGTGAATCATAAGCGCGGTT |
| ***cry2*_pMP7 R** | CAGACAATTGACGGCTCTAGAAACTCGGTAATAGGCAGAA |
| **pMP7_bb_Gibson F** | TCTAGAGCCGTCAATTGTCTG |

| | |
|---|---|
| **pMP7_bb_Gibson R** | GAATTCGATATCAAGCTTATCG |
| **RT-qPCR** | |
| **gyrA_rt_F** | GAGCCAAAGTTACCTTGGCC |
| **gyrA_rt_R** | AATGTGCTGGGCAACGACTG |
| **aspA_rt_F** | TCTGCGGCGAATGTAATGGT |
| **aspA_rt_R** | TCCATCATGCCAGCGAAAGT |
| **dcuA_rt_F** | TGAAATCGGTGAAGCGGTGA |
| **dcuA_rt_R** | ACCGCATTTTCCACTTTGCC |
| **rimO_rt_F** | TGATTGTGTACCGGATGCGG |
| **rimO_rt_R** | TTAAAGTCTGGTCGTGCGCC |
| **rstA_rt_F** | ATCTGGGCGCCTCTTCCTAA |
| **rstA_rt_R** | CACCTCCAAGCGTTCCATCA |
| **rstC_rt_F** | GTTCAGGCGCTTATACAGACGA |
| **rstC_rt_R** | GTTGCGGATTTAGGCTTGGTG |
| **vc02680_rt_F** | CAAAACCCAATCCCACTGCG |
| **vc02680_rt_R** | CGTGGACTTGCAGAACTTTCG |
| **vca02766_rt_F** | TCCGACGTTAGACAAGTGGC |
| **vca02766_rt_R** | CCTTTCATCACCGGAGCACT |

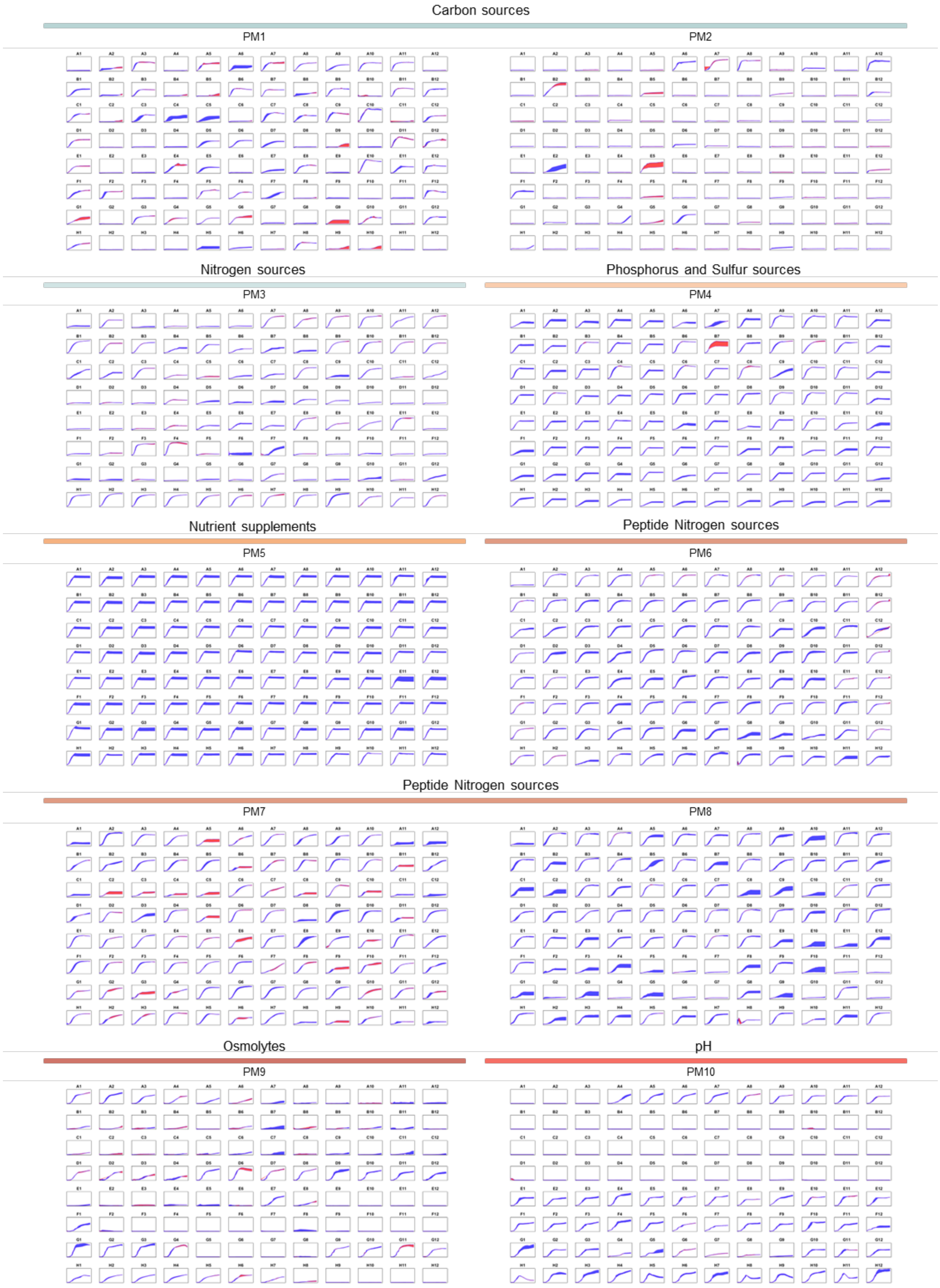
Carbon sources
PM1
PM2
Nitrogen sources
PM3
Phosphorus and Sulfur sources
PM4
Nutrient supplements
PM5
Peptide Nitrogen sources
PM6
Peptide Nitrogen sources
PM7
PM8
Osmolytes
PM9
pH
PM10

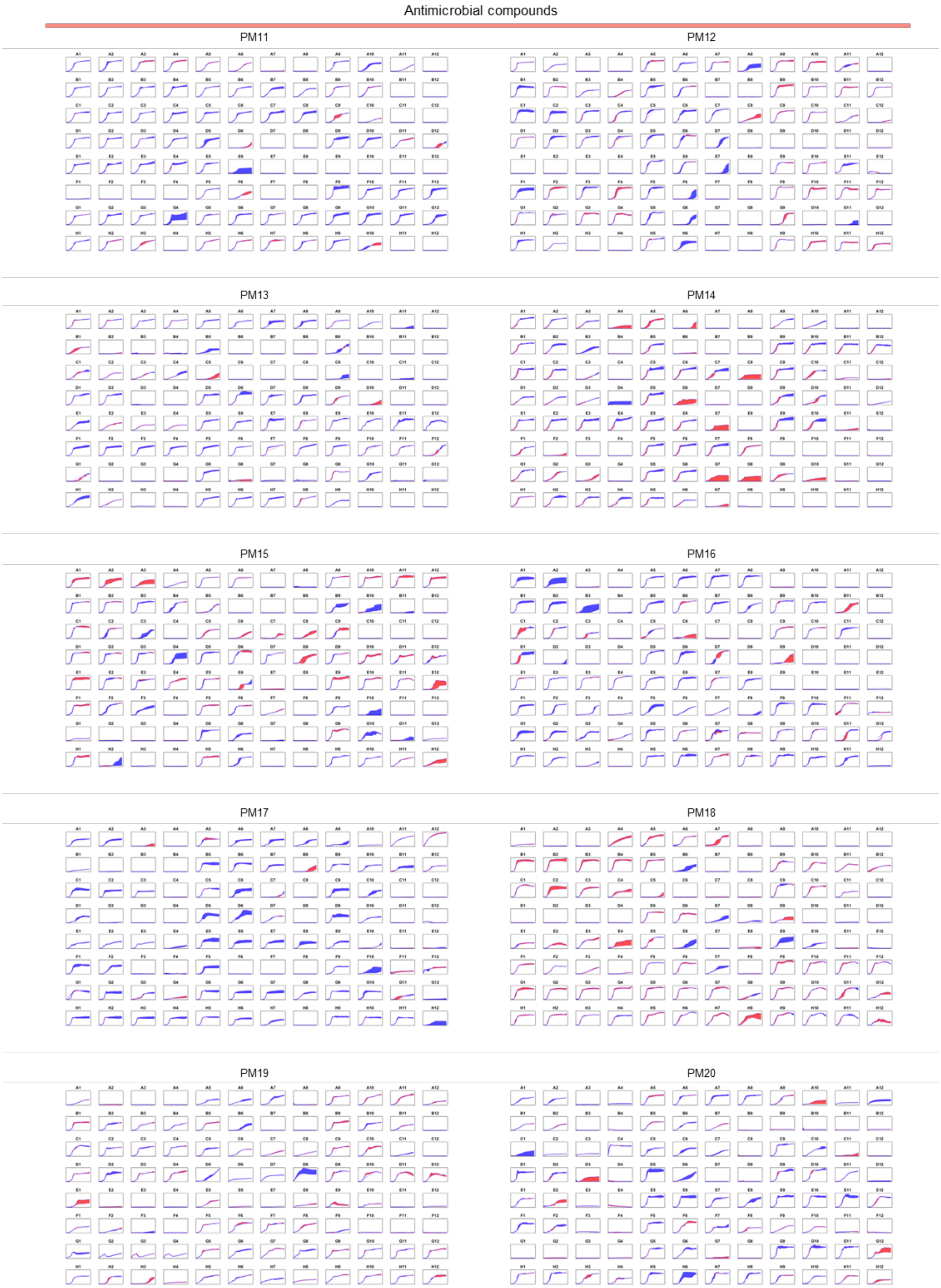
Antimicrobial compounds
PM11
PM12
PM13
PM14
PM15
PM16
PM17
PM18
PM19
PM20

**Supplementary Figure S1.** Growth curves representation of *V. cholerae* WT compared to the ΔSI mutant in all Biolog Phenotype microarrays plates (PM1-20). The mean growth curve from two independent biological replicates of the WT strain is graphically overlaid with the mean growth curve of the mutant strain calculated from two independent replicates. If the ΔSI mutant has an impaired growth in comparison with the WT, the overlapped area in the corresponding well in the graph will be blue, whereas if the ΔSI mutant has a growth improvement in comparison with the WT, the overlapped area in the corresponding well will be red.

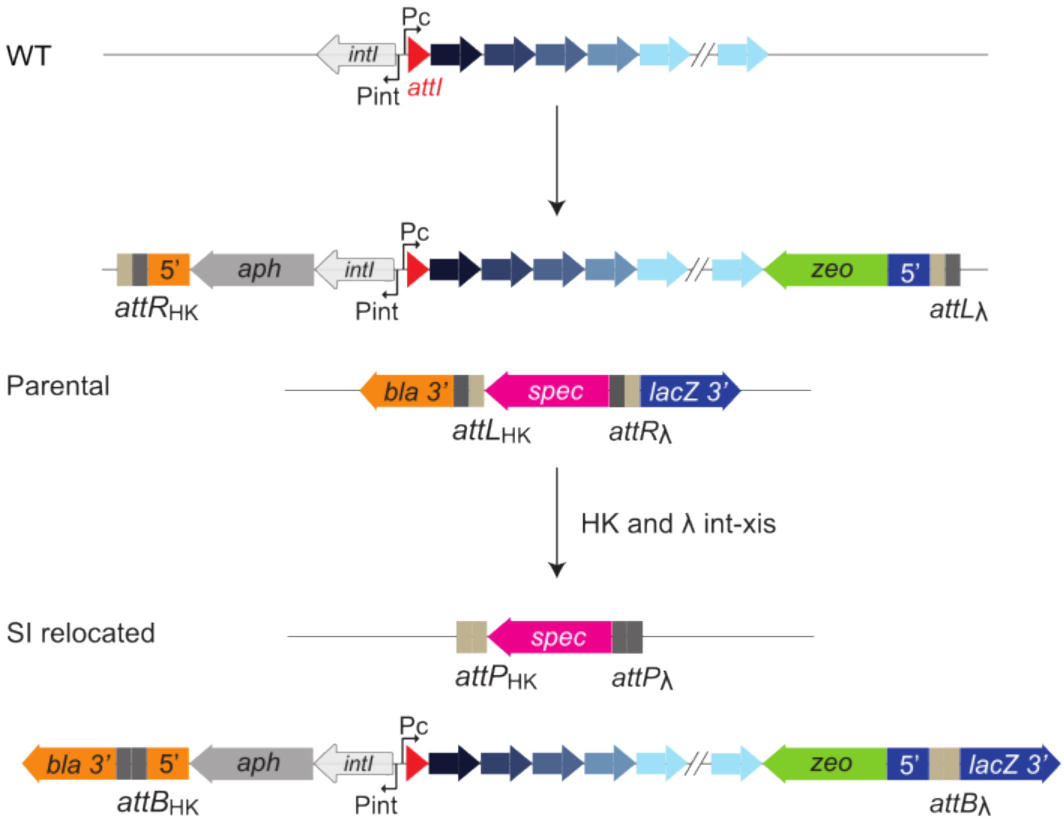


**Supplementary Figure S2.** Schematic representation of the relocation of the *V. cholerae* SI from chromosome 2 to chromosome 1. The relocation is based on the recombination of two bacteriophage attachment sites, named *attL* and *attR*. These sites are associated with fragments of a genetic marker that becomes reconstituted when they recombine (*bla* and *lacZ*), allowing for the selection of those clones where the SI has been relocated.
